# Spatial patterning of transcriptional and regulatory programs in the primate subcortex

**DOI:** 10.1101/2025.11.22.689869

**Authors:** Shu Dan, Meghan A. Turner, Michael DeBerardine, Lakme Caceres, Victor E. Nieto-Caballero, Jessica Schembri, Kirsten Levandowski, Christopher Cardenas, Ning Wang, Chang Li, Soumyadeep Basu, Hongyu Zheng, Xinhao Liu, Thomas Höllt, Jessica Feliciano, Qiangge Zhang, Reilly Nakamoto, Delissa A. McMillen, Naomi Martin, Nasmil Valera Cuevas, Paul Olsen, Josh Nagra, Jazmin Campos, Marshall M VanNess, Jack Waters, Shea Ransford, Zoe Juneau, Sam Hastings, Stuard Barta, Augustin Ruiz, Jeanelle Ariza, Benjamin J. Raphael, Hongkui Zeng, Ed S. Lein, Boudewijn Lelieveldt, Guoping Feng, Brian Long, Fenna M. Krienen

## Abstract

Mammalian brain cell identity is shaped by intrinsic factors and external context. We present a spatially resolved transcriptomic and gene regulatory atlas of cell types across all subcortical regions in a primate, the common marmoset. Dense sampling and cross-species integration revealed spatially precise neuronal assemblies, including in complex midbrain and diencephalic structures. Chromatin accessibility and transcriptional identity are spatially tuned within and across subcortical structures; spatial gradients within hippocampal subfields are orchestrated by graded transcription factors acting through graded enhancers. The primate-expanded thalamic GABAergic population shares transcriptional and regulatory syntax with conserved midbrain populations, reflecting an evolutionary adaptation compared with rodents. Similar regional expression across cell types can arise by distinct regulatory architectures, as for telencephalic astrocytes and neurons. Conversely, distant cell types can share regulatory programs despite divergent identities: striatal GABAergic medium spiny neurons and telencephalic glutamatergic neurons share a postsynaptic regulatory program despite divergent lineage, region, and neurotransmitter identity.

## Introduction

Single cell genomics studies have transformed our understanding of cellular diversity; the brain is a particularly fertile ground for these efforts. Recent data indicate that thousands of transcriptomically distinct cell populations exist in the adult mouse brain^1,2^, which has the most comprehensively characterized brain cell taxonomy of any mammal. While the mouse remains foundational for understanding the mammalian brain, the extent to which mouse cell types and their spatial organization generalize to primates remains an open question.

The human brain is ∼3,000-fold larger by mass than the mouse brain and contains more than 86 billion neurons. This scale makes comprehensive study a formidable challenge. The most extensive human brain atlas to date^3^ used snRNA-seq to sample over 100 anatomical locations of the adult human postmortem brain, but anatomical assignments are relatively coarse and based on microdissection labels. More generally, the size of the human brain precludes complete spatial transcriptomic coverage with current methods. These limitations obscure rare cell types, the precise anatomical localization of individual populations, and complex spatial topographies such as gradients and discontinuities. In turn, incomplete sampling complicates cross-species comparisons, particularly as cell migration is a driver of evolutionary change^4–7^.

The marmoset *Callithrix jacchus* presents a compelling solution to the issue of scale. The marmoset is a New World monkey whose brain weighs only 8 grams, comparable to that of a rat^8,9^. Despite its evolutionary distance to humans (35-40 million years to great apes), the adult marmoset brain retains primate-specific anatomical^10–14^, connectivity^15–17^, and cell type specializations^18–21^. With its compact brain and complete anatomical accessibility via current single-cell and spatial transcriptomic platforms, the marmoset is uniquely suited to fill in the gaps in our understanding between mouse and primate cell type diversity.

While transcriptomic atlases of the mouse brain are increasingly detailed, the gene regulatory architecture supporting this diversity remains largely unexplored at brain-wide scale. Single-region studies link chromatin accessibility and cis-regulatory elements to cell identity and spatial context^22–26^, but no spatially-resolved regulatory atlas spans numerous brain regions in a mammalian species. Such an atlas would inform how cell lineage and regional context shape cell identity and the regulatory logic of spatial gradients, expand the search space for enhancer-based genetic tools across species^20,27–31^, and provide a foundation for interpreting noncoding disease variants.

Here, as part of the BRAIN Initiative Cell Atlas Network (BICAN, RRID:SCR_022794), we generated a hierarchical cell taxonomy based on anatomically comprehensive multiomic (snRNA-seq and snATAC-seq) sampling of the marmoset subcortex. In parallel, we performed spatial transcriptomic profiling of a marmoset cerebral hemisphere, enabling anatomical localization of cell populations at each level of the taxonomy. All three modalities can be explored interactively through our Cytosplore Viewer (https://viewer.cytosplore.org/). With this atlas, we characterize gene regulatory networks in both neurons and non-neurons that distinguish cell types and brain regions, dissect convergent and divergent regulatory programs across cell types, characterize spatial gene expression gradients in the hippocampal formation, and investigate the evolutionary origins of primate-expanded thalamic GABAergic neurons. Our study analyzes the complete subcortex in marmoset, while companion BICAN papers focus on basal ganglia cell types across human and non-human primates ^32,33^.

## Results

### Generation of the marmoset subcortical cell atlas

The marmoset subcortical cell atlas (MSCA) encompasses the non-isocortical telencephalon, including the hippocampal formation (HiF), olfactory areas (OlfC), and cerebral nuclei (CNU), as well as the diencephalon (thalamus and hypothalamus), midbrain, and major hindbrain structures (excluding cerebellar cortex). We integrated the MSCA with existing human, non-human primate, and mouse brain atlases to support interoperable annotations and cross-species comparisons. MSCA data that mapped to the basal ganglia nuclei (BG) structures were also used in the companion study describing a spatial atlas of marmoset, macaque and human BG^33^ as well as in a consensus cross-species multiomic BG taxonomy^32^.

#### An anatomically comprehensive cell type taxonomy of the marmoset subcortex

The subcortex of four marmoset donors was tiled into contiguous dissections covering the entire subcortex (**Fig 1A**, **Table S1**, STAR Methods). Dissociated nuclei from each tile were fluorescence-activated nuclei sorting (FANS)-sorted to enrich for neurons and deplete oligodendrocyte lineage populations following the protocol in Johansen, Fu, Schmitz et al^32^. Sorted fractions were repooled to target proportions (70% NeuN+, 10% Olig2+, 20% NeuN-/Olig2-), then profiled by 10X Multiome. In total, we sequenced 1,299,000 nuclei from 76 unique tiles across the four donors, targeting a sequencing depth of 120,000 reads/cell.

**Figure 1.**
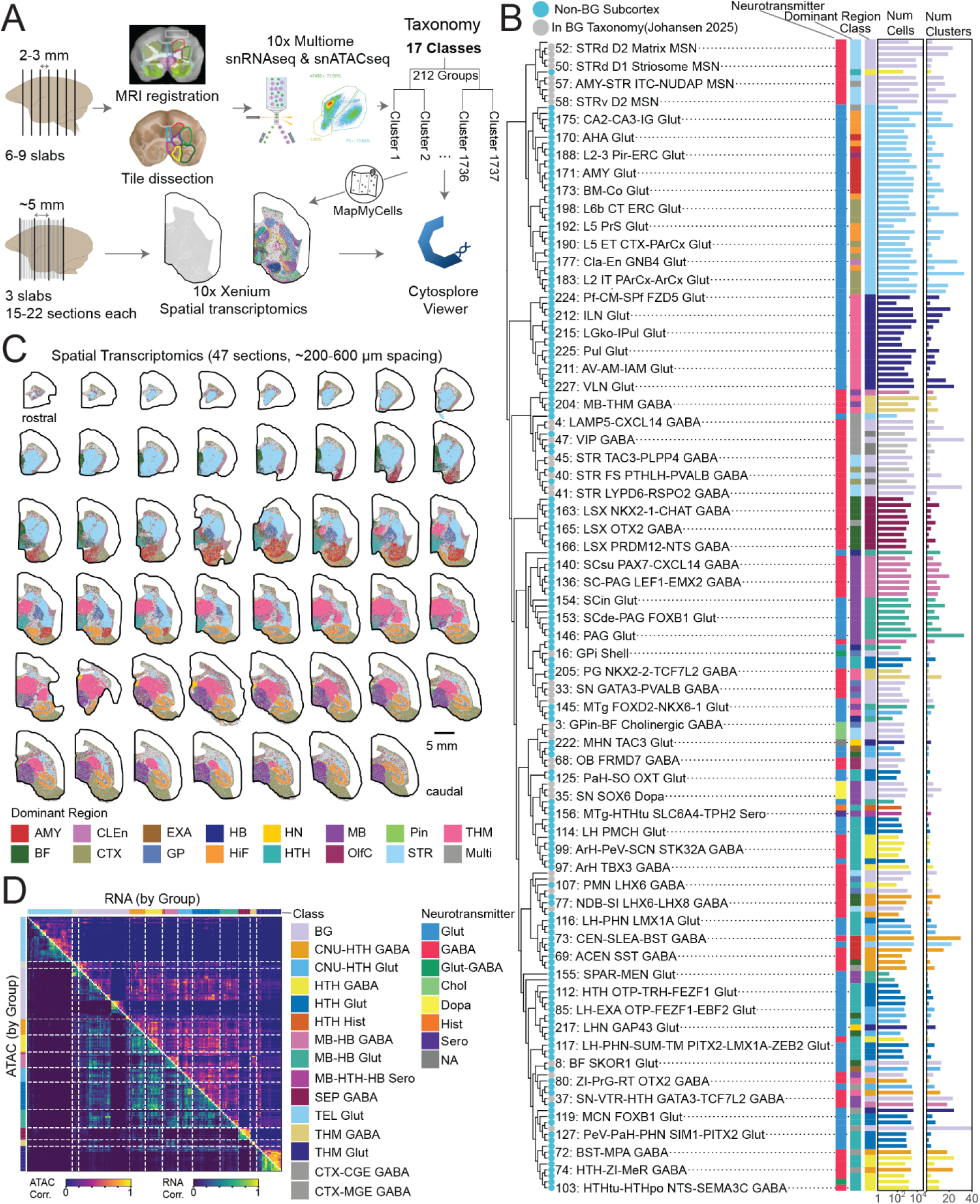
Spatial, transcriptomic, and regulatory atlas of the marmoset subcortex. **(A)** Schematic of sampling and analysis workflows. Nuclei from MRI registration-guided dissection tiles were FANS-sorted for 10X Multiome, yielding a three-level taxonomy (Class, Group, Cluster). Xenium spatial transcriptomics data were mapped to the taxonomy to obtain spatially-resolved Cluster identities. Cytosplore Viewer links multiomics and spatial transcriptomics data for interactive data exploration. **(B)** 194 neuronal transcriptomic Groups hierarchically clustered based on the expression of top 4000 HVGs. Group IDs and names labelled every third branch. Groups included in the companion cross-species consensus BG taxonomy^32^ are indicated before Group names. Color tiles (left to right) show the neurotransmitter, dominant region, and Class. Bar plots indicate cell and Cluster counts per Group. **(C)** 47 Xenium spatial sections (rostral to caudal); each cell colored by the dominant region of its assigned Cluster. Scale bar, 5 mm. **(D)** Pairwise Spearman correlation of transcriptomic and chromatin accessibility similarity between all Groups; ordered as in (B).

After quality control (STAR Methods, **Fig S1**), 678,917 nuclei remained. Batch effects were corrected by scVI^34^ across donors, and iterative clustering^35^ in the donor-aligned latent space resolved 1737 transcriptomic “Clusters”. We organized Clusters into a three-level hierarchy by aggregating transcriptomically and spatially similar Clusters into 212 Groups and 17 Classes (Methods, **Fig 1A-B**). Groups are generally named for the anatomical structures where the majority of cells are found. Clusters belonging to BG inherited their Class and Group labels from the BG Cross-Species Consensus Taxonomy^32^.

We hierarchically clustered Group-level pseudobulked “metacells” using the top 4000 highly variable genes (HVGs) (STAR Methods, **Fig 1B**). Within neurons, clades of excitatory telencephalic (cortex (CTX), amygdala (AMY), and HiF) Groups align with major anatomical boundaries. Notably, the striatal medium spiny neurons (MSNs), although inhibitory, clustered with excitatory telencephalic Groups rather than other telencephalic GABAergic populations, consistent with prior mouse and primate reports^20,36,37^.

In the diencephalon, excitatory thalamus (THM) Groups formed a distinct clade, with inhibitory thalamic Groups more like inhibitory Groups from other structures. By contrast, hypothalamic Groups intermingled regardless of neurotransmitter identity. Other major branches reflected neurotransmitter (e.g. dopaminergic and cholinergic Groups) or anatomical (e.g. lateral septum (LSX), superior colliculus (SC)) identity.

#### Spatially resolving subcortical cell type diversity and mapping performance

Spatial transcriptomic data (10X Xenium) were collected from one hemisphere at ∼200 µm section spacing across three 5-mm slabs (STAR Methods; **Fig 1C**). We designed a 300-gene panel (**Table S2**) from previously described cell-type markers, differentially expressed genes within caudate and putamen^12,38^, and variable genes from prior marmoset BG and thalamic snRNA-seq^20^. Spatial cells were mapped to the MSCA taxonomy using correlation-based mapping (MapMyCells^39^ RRID:SCR_024672) at Cluster level, with Group and Class labels inherited.

To evaluate panel performance, we mapped multiome RNA back to its own taxonomy using only the 300 spatial panel genes (self-mapping; STAR Methods, **Fig S2**). This recovered the Dominant Region annotation for 97.5% of neurons in localized Clusters (93% of all taxonomy neurons) (**Fig S2C**). The panel performed well for its size and was balanced across the taxonomy (**Fig S2D-F**).

In self-mapping, ∼99% of cells receive a correct Class assignment and >90% of cells receive a correct Group assignment; rates varied by cell type **(Fig S2G-I)**. Cluster-level accuracy varied across the taxonomy (60% overall, with confusions concentrated amongst transcriptomically similar neighbors in the same structure **(Fig S2J; Table S3)**. After pseudobulking spatial data by assigned Cluster, 71.5% of Clusters were most correlated with their multiome counterpart, and 87.0% fell within the top two matches. Class- and Group-level cell abundances were broadly consistent between modalities (**Fig S2K-L**). Per-Cluster and per-Group performance is reported in **Table S3**.

#### Spatial mapping reveals anatomically coherent cell type organization

Spatial mapping revealed detailed localization of both neuronal and non-neuronal populations and allowed anatomical annotation of transcriptomic Groups; specific examples are shown in **Fig S3-S5**. Within the amygdala, Groups mapped to individual nuclei via discrete markers and Clusters resolved finer positions through combinatorial and graded expression (**Fig S3**). Comparable anatomical specificity was observed in thalamic glutamatergic populations (**Fig S4**) and non-neuronal populations of neuroglia and mesodermal lineages (**Fig S5**).

#### Integrating cross-species references into marmoset subcortical cell type annotations

To further support our cell type annotations, we cross-referenced our taxonomy against additional publicly available mouse^1^ and human^3^ whole brain atlases through MapMyCells^39^ (STAR Methods, **Tables S4-5**). We also integrated regional atlases for the telecephalon^20^, AMY^40^, HTH^41,42^, LSX^43^, HiF^44^, and lateral geniculate nucleus (LG)^45^ using scVI^34^ (STAR Methods, **Fig S6A-B**, **Tables S4-5**). Despite differences in species and assay modality, MSCA cell type divisions and spatial annotations were largely concordant with the transcriptomic groupings and anatomical dissections in these reference resources.

Companion BICAN papers use MSCA BG cells in a cross-species spatial BG atlas^33^ and consensus cross-species multiomic BG taxonomy^32^. The contiguous sampling of the MSCA beyond BG structures reveals relative similarities of cell populations within and between BG and non-BG structures (**Fig 1B**). For example, MSCA mapping shows that the evolutionarily divergent striatal TAC3+ interneuron population^4,5,18,20,46^ includes subtypes distributed outside the striatum – in septum, ventral pallidum, and HTH – indicating greater molecular and anatomical diversity than previously appreciated (**Fig S6C**).

#### Chromatin accessibility recapitulates the transcriptional hierarchy

Gene expression is controlled by a vast diversity of cis-regulatory elements (CREs, including enhancers). Millions of putative enhancers decorate mammalian genomes^47,48^, any one gene may have dozens of enhancers, and a given cell type may have thousands of enhancers that are differentially accessible compared to other cell types. From Group-level ATAC peaks (SnapATAC2^49^; STAR Methods), we identified ∼462,000 CREs robustly accessible in at least one Group. Topic modeling (pycisTopic^50^) grouped peaks into modules and yielded a low-dimensional accessibility embedding. The ATAC-based embedding closely mirrored the RNA-based embedding, suggesting that the regulatory syntax captured by the topics recapitulates the transcriptional hierarchy (**Fig S7A-B**). Group-level marker genes showed concomitant open chromatin along their gene bodies in the ATAC data (**Fig S7C**), indicating that Group-level differences were preserved in the ATAC data.

Hierarchical clustering of pycisTopic ATAC loadings across Groups (**Fig 1D**, bottom triangle; **Fig S8**) closely matched Group-level RNA correlations (based on 4000 HVGs, **Fig 1D**, top triangle; **Fig S8**) (Mantel test with 5000 permutations, r=0.88, p<0.001). This indicates that gene regulatory and gene expression landscapes converge on a globally concordant view of cell identity (**Fig S7D**).

#### Cross-species mapping resolves heterogenous subcortical populations

We leveraged the MSCA to provide greater resolution to the most anatomically complete human brain cell taxonomy^3^, which sampled 100 locations by snRNA-seq but lacked precise spatial information. The human cell taxonomy contains 31 Superclusters, 461 Clusters, and 3313 Subclusters. We used MapMyCells to transfer human Supercluster and Cluster labels onto marmoset cells (**Fig 2A**, STAR Methods). On average, 98% of cells in each MSCA Cluster mapped to the same Supercluster. Homologous MSCA Groups were identified for each of the 20 neuronal Superclusters (excluding the human cerebellar inhibitory Supercluster due absent cerebellar sampling) (**Fig S9**). The “*Splatter*” Supercluster mapped to the most MSCA Groups (n=117), Clusters (n=974), and cells (n=126941) (**Fig. 2B**; **Fig. S9**), spanning nearly a dozen anatomical regions (**Fig. S10** inset).

**Figure 2.**
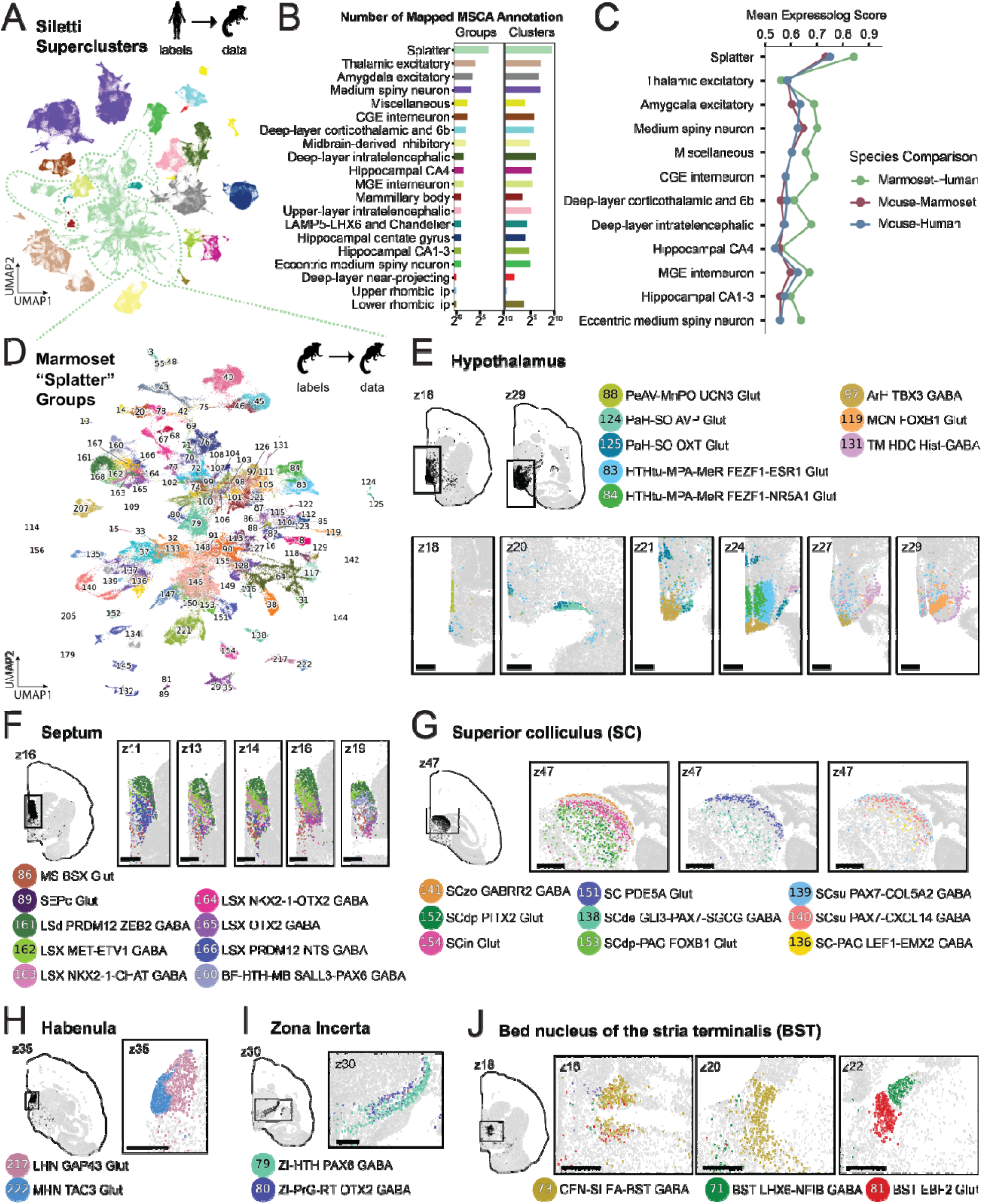
Cross-species spatial mapping of heterogeneous neuron populations. **(A)** UMAP of marmoset subcortical neurons colored by predicted human Supercluster identity from Siletti et al^3^. **(B)** Number of MSCA Groups and Clusters mapped per human Supercluster. **(C)** Mean expressolog scores of homologous clusters across human, marmoset and mouse within each human Supercluster; higher scores indicate stronger transcriptomic similarity. **(D)** UMAP of marmoset neurons mapped to the *Splatter* Supercluster, colored by MSCA Group; Group IDs overlaid (**Fig S10** and **Table S4** for full names). **(E-J)** Representative spatial z-planes showing MSCA Group localization; whole-hemisphere panels provide anatomical context. Scale bars, 1 mm. **(E)** HTH structure Groups. **(F)** Medial and lateral septum Groups. **(G)** Layers of SC. **(H)** Medial and lateral habenula. **(I)** ZI subdivisions. **(J)** BST subdivisions.

To test conservation across mammals, we defined homologous neuronal cell types as mouse^1^ or marmoset Clusters for which >70% of cells mapped to a given human Cluster, yielding 114 of 382 human clusters with conserved counterparts in both species. Pairwise cross-species expression similarity, quantified by expressolog^51^, confirmed that marmoset cells are more similar to those in human than to mouse (**Fig 2C**). Notably, the *Splatter* Supercluster is both the most broadly mapped Supercluster and the most conserved across all species pairs, despite its transcriptional heterogeneity.

The *Splatter* Supercluster was named for its complex topography in low dimensional embeddings. Siletti et al.^3^ attributed this to neurons from diverse regions (basal forebrain, hypothalamus, thalamus, midbrain, pons, medulla) clustering together due to under-sampling; our dense sampling and spatial data now resolve their anatomical origins (**Fig 2D, Fig S10**). One of the largest *Splatter* subpopulations localized to the marmoset hypothalamus (HTH) (**Fig 2E**), which contains many small and functionally distinct neuronal populations^41,42,52,53^. Consistent with integration of mouse and human “Hypomap” atlases^41,42^, many marmoset Groups are localized to specific hypothalamic nuclei including the arcuate nucleus (ArH) and magnocellular nucleus (MCN) (**Fig 2E, Fig S6B**). Other Groups resolved subdivisions within nuclei, such as the distinct medial and lateral Groups within the ventromedial hypothalamus of the tuberal nucleus (HTHtu) (**Fig 2E**). The mammillary body (MM) – its own Supercluster in the Siletti human atlas – is subdivided by marmoset Clusters into six spatially distinct partitions along medial-lateral and rostral-caudal axes (**Fig S11A**).

Where regional reference annotations were sparse (basal forebrain and midbrain), we leveraged the spatial transcriptomics data directly. In the basal forebrain, the septum is patterned by seven GABAergic and two glutamatergic Groups, anatomically anchoring homologous cell types from an integrated human septum atlas^43^ (**Fig 2E, Fig S6B**). Groups mapping to the midbrain superior colliculus (SC) highlighted its internal laminar organization (**Fig 2G**). Distinct Groups partitioned the medial (MH) and lateral habenula (LH) (**Fig 2H**) and the zona incerta (ZI) (**Fig 2I**); in the extended amygdala, three Groups delineated the rostral-caudal axis of the bed nucleus of the stria terminalis (BST) (**Fig 2J**). Further spatial subdivisions are present at the Cluster level within LH/MH (**Fig S11B**), ZI (**Fig S11C**), and BST (**Fig S11D**).

Collectively, these mappings resolve the human *Splatter* Supercluster into hundreds of anatomically discrete, evolutionarily conserved populations distributed across the hypothalamus, basal forebrain, midbrain, habenula, zona incerta, and extended amygdala. It reframes what had appeared as a single heterogeneous group in human data^3^ as a spatially organized collection of conserved subcortical cell types.

### Gene regulatory programs across the primate subcortex

A key benefit of paired RNA and ATAC multiome is the ability to identify enhancers that link TFs to target genes, forming gene regulatory networks (GRNs). We inferred GRNs using SCENIC+^54^, which integrates chromatin accessibility, gene expression, and transcription factor binding site (TFBS) motif analysis to identify eRegulons that link TFs to binding sites and target genes (**Fig 3A**). ATAC peaks within eRegulons are candidate cis-regulatory elements (cCREs), predominantly enhancers with fewer repressors. We identified 550 neuronal and 299 non-neuronal eRegulons (STAR Methods, **Table S6**).

**Figure 3.**
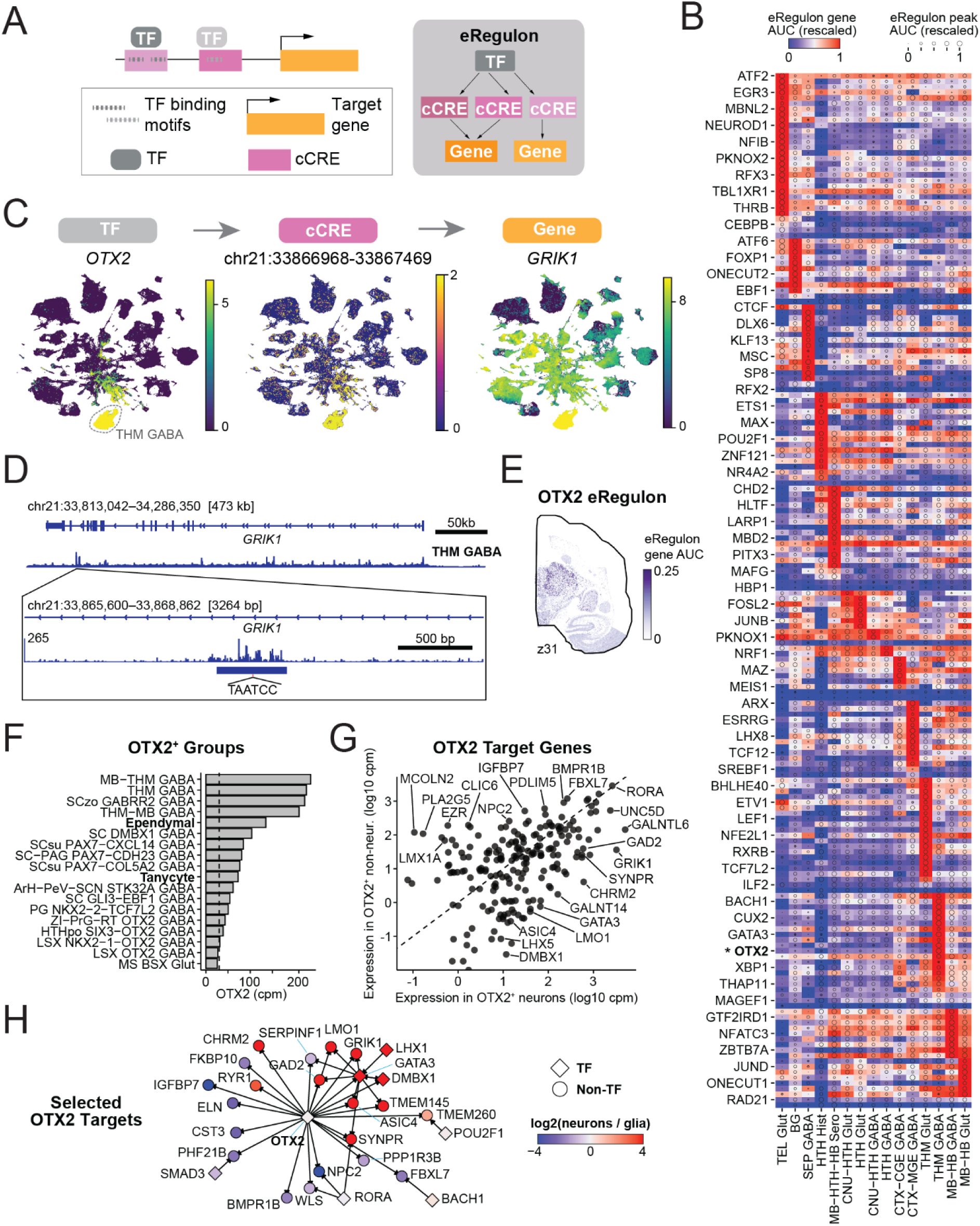
Gene regulatory networks across primate subcortex. **(A)** Schematic eRegulon components (adapted from Bravo González-Blas et al.^54^). **(B)** Direct activator (“+/+”) eRegulons for each neuronal Class; every third eRegulon labelled. *OTX2* eRegulon denoted by asterisk. **(C)** UMAP plots for an example eRegulon triplet; peak accessibility and target gene expression correlated with *OTX2* expression (*r*=0.77 and 0.57, respectively). **(D)** Within Class THM GABA, ATAC-seq insertion events near the peak and target gene from (B); the intronic peak contains a canonical *OTX2* motif. € Representative spatial section showing high *OTX2* eRegulon activity in the thalamus. **(F)** Top Groups by *OTX2* expression; non-neuron groups in bold. **(G)** Expression of *OTX2* target genes in neuronal vs. non-neuronal Groups, with select shared and differentially enriched genes highlighted. **(H)** Network plot for selected neuron- or non-neuron-specific OTX2 targets (STAR Methods).

#### Conserved transcription factor OTX2 illustrates cell-type-specific regulatory logic

While some GRNs show broad activity, most are active in specific classes of neurons or non-neurons (**Fig 3B**, **Fig S12A**). One striking case is the *OTX2* eRegulon, highly active in Group “THM GABA” (**Fig 3B**). *Otx2* is a developmental regulator of the telencephalon, diencephalon, and mesencephalon and specifies the midbrain-hindbrain boundary^55–57^. In neurons, *OTX2* has 142 putative direct targets including *GRIK1* (GluK1 kainate receptor subunit) (**Fig 3C**). SCENIC+ predicts *OTX2* activity at multiple peaks associated with *GRIK1* expression, including an intronic peak with a consensus *OTX2* motif whose accessibility correlates with both *OTX2* and *GRIK1* expression (**Fig 3D)**.

Projection of eRegulon activity onto spatial data confirmed that *GRIK1* and other neuronal *OTX2* targets are concentrated in the thalamus (**Fig 3E**). Other Groups with high *OTX2* expression include midbrain GABAergic types (especially in SC) as well as ependymal glia and tanycytes (**Fig 3F**), paralleling the adult mouse brain^1^. We also detected a non-neuronal *OTX2* eRegulon, with 69 additional targets. Many *OTX2* targets show cell-type-divergent expression, including the TFs *DMBX1* and *GATA3* (neuron-specific), and *LMX1B* and *PDLIM5* (non-neuron-specific) (**Fig 3G**).

How can *OTX2* be expressed in two cell populations yet have different targets in each? We hypothesized that cell-type-specific co-TFs may be acting with *OTX2* at cell-type-specific enhancers. cCREs for cell-type-specific targets show cell-type-specific accessibility (**Fig S12B-C**), consistent with co-TF-gated accessibility. Searching for TFs co-enriched in OTX2 cCREs (n=152 neuronal, n=46 non-neuronal) identified strong DMBX1 and GATA3 enrichment in neurons (n=36 and n=20 shared gene targets, respectively; **Fig S12D**). DMBX1 and OTX2 share a motif in their shared peaks, and they physically interact^58^, supporting DMBX1 as a potential neuronal co-factor – though this was supported at only a minority of neuronal OTX2 sites. In non-neurons, NFIA was most co-enriched, albeit with few shared targets (n=4).

#### Convergent regional expression in neurons and astrocytes arises through distinct regulatory programs

The OTX2 example shows that a single TF can direct distinct targets across cell types. We next examined the converse: can the same regionally-biased gene be expressed convergently in different cell types through distinct regulatory architectures? Astrocytes and neurons provide a natural testbed because both are regionally heterogeneous in the mammalian brain^59–61^. We identified three major astrocyte subdivisions: Astro-TE NN, Astro-NT NN, and Astro-WM NN Groups (**Fig 4A**; **Fig S12E**), paralleling the three regional identities reported in mouse^1^ and corresponding to the primate-conserved GM-STR, GM-exSTR, and WM subgroups described in Fu et al^62^. Principal component analysis (PCA) on pseudobulked astrocyte Clusters (**Fig S12F**) separated Astro-NT NN Clusters from Astro-TE NN and Astro-WM NN Clusters along PC1. Top PC1 genes included *FOXG1* – a known telencephalic regulator in both astrocytes and neurons^63,64^ that we revisit in later analyses – and *MOXD1,* a copper dependent monooxygenase, which was enriched in Astro-TE NN and Astro-WM NN Groups (**Fig 4B**). *MOXD1* showed the same regional signature in neurons: enriched in telencephalic neurons, particularly in amygdala and septum (though at lower levels than in Astro-TE and Astro-WM), and largely absent from non-telencephalic neurons (**Fig 4C**).

**Figure 4.**
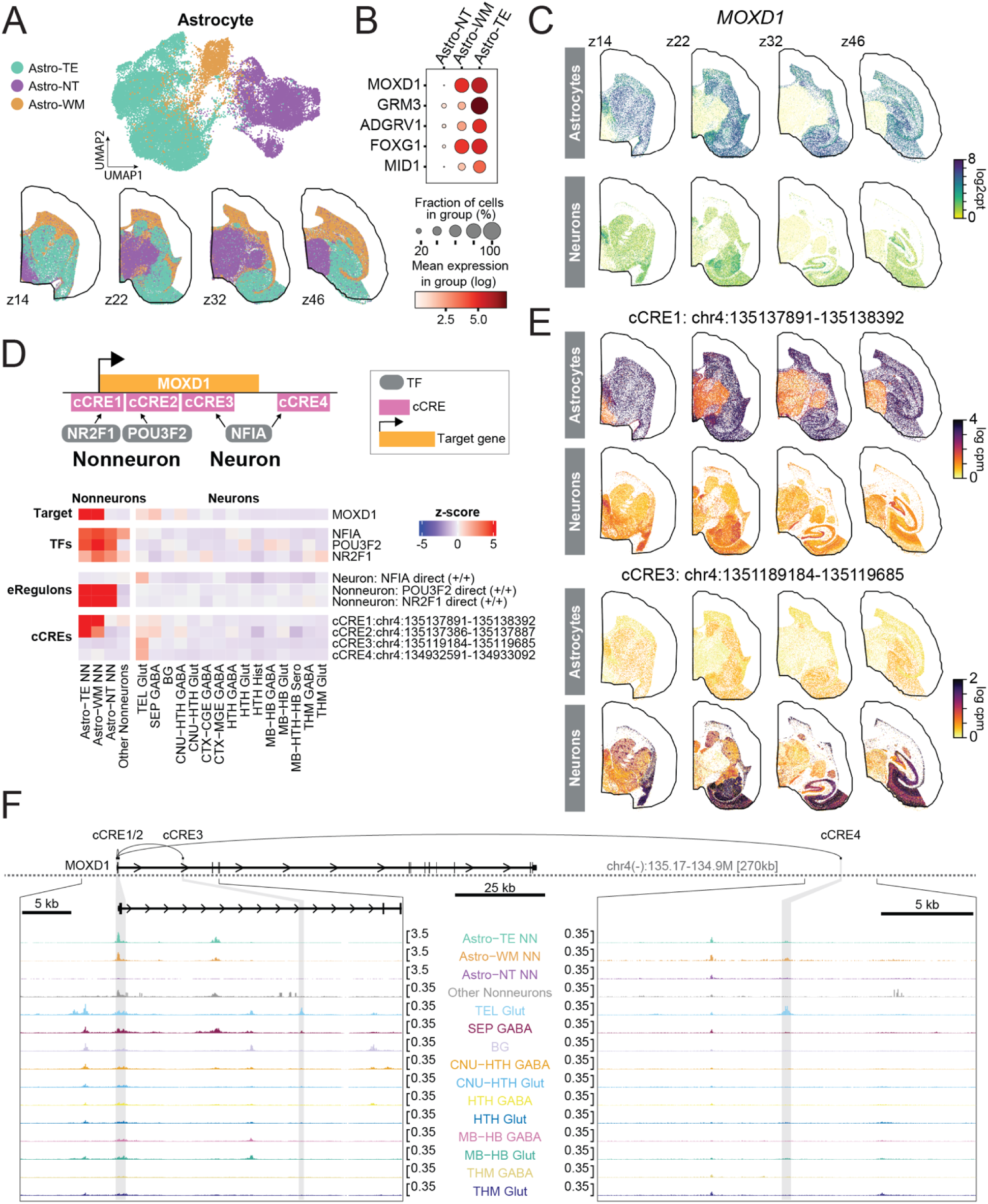
Regional zonation of astrocytes. **(A)** Regional zonation of the three astrocyte Groups (Astro-NT NN, Astro-TE NN, Astro-WM NN); UMAP embedding (top) and representative spatial sections (bottom) colored by Group. **(B)** Dot plot of top PC1 loading gene expression across astrocyte Groups. **(C)** *MOXD1* expression (log2cpt) in all astrocytes (top) and all neurons (bottom) on representative spatial sections. **(D)** *MOXD1*-related eRegulons in non-neurons and neurons. Top: Schematic summarizing the TFs and cCREs driving *MOXD1* expression (see Results). Bottom: z-scored heatmaps of each component. **(E)** Representative spatial sections showing Cluster-average chromatin accessibility (log2cpm) for an astrocyte-specific cCRE(1) and a neuron-specific cCRE(3). **(F)** Browser tracks of *MOXD1*-related cCREs grouped by Class.

*MOXD1* was linked to distinct regulatory programs in neurons and astrocytes. In astrocytes, two eRegulons driven by *NR2F1* and *POU3F2* were active across all astrocyte Groups, with cell-type specificity achieved through selective accessibility of promoter cCREs (cCRE1, cCRE2) only in Astro-TE and Astro-WM. In neurons, *MOXD1* was a target of an *NFIA*-driven eRegulon acting at intronic (cCRE3) and intergenic (cCRE4) sites (**Fig 4D-F**). Thus, convergent regional expression of *MOXD1* in telencephalic neurons and astrocytes arises through distinct regulatory architectures.

Cross-species comparison revealed species differences: mouse astrocytes lack *Moxd1* while telencephalic neurons retain it (**Fig S12H**); human astrocytes show telencephalic enrichment like marmoset (**Fig S12I**). This pattern is consistent with prior developmental studies showing that *MOXD1* is enriched in the glial progenitors of human outer radial glia during cortical development, whereas *Moxd1* is not detected in comparable mouse radial glial populations^65,66^, raising the possibility that telencephalic and white-matter astrocyte *MOXD1* expression reflects retention of a primate-associated developmental program.

#### Developmental origin and regional context shape neuronal regulatory identity

We next examined how regulatory programs track anatomical location and developmental origin in neurons. Some progenitors – notably those that produce telencephalic GABAergic neurons – can disperse lineage-related daughters across regions^67–70^. This provides opportunities for regulatory divergence with shared lineage, or for regulatory convergence across distinct lineages.

To dissect these contributions, we regrouped regionally localized neuron Clusters by anatomical region (from spatial mapping) and major neurotransmitter, subdividing ganglionic eminence (GE)-derived neurons (GABAergic or cholinergic) by progenitor origin (MGE, CGE, or LGE), and separating MSNs from other GABAergic neurons. We then compared eRegulon usage across groups (**Fig 5, Fig S13**).

**Figure 5.**
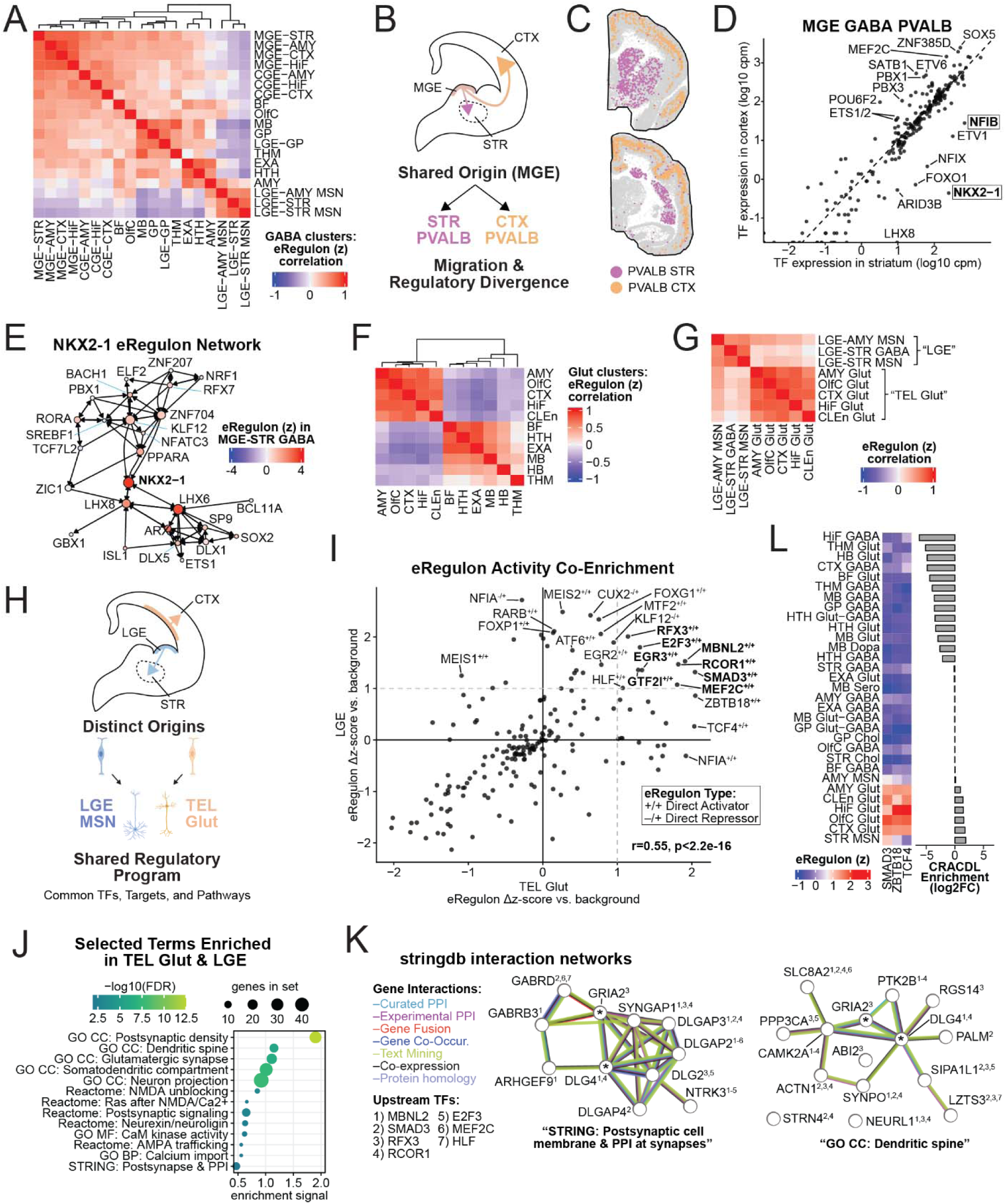
Regional dynamics of gene regulatory network activity. **(A)** GABAergic Cluster-level eRegulon scores averaged by region, with MSNs and GE-derived clusters separated; Pearson correlations shown. **(B)** Schematic: MGE-derived GABAergic neurons migrate to both cortex and striatum. **(C)** Spatial mapping of STR- and CTX-localized PVALB interneurons. **(D)** TF expression comparison in STR- vs CTX-localized PVALB clusters; similar NKX2-1 and NFIB patterns seen in SST neurons (**Fig S13B**). **I** TF-TF eRegulon network centered on NKX2-1, colored by eRegulon activity averaged across MGE-STR GABA clusters. **(F)** Glutamatergic clusters compared as in (A). **(G)** LGE-derived GABAergic neurons and telencephalic glutamatergic neurons are highly similar and distinct from other neurons. (see also **Fig S13A**). **(H)** Schematic of regulatory convergence between TEL Glut and LGE GABA. **(I)** eRegulon activity enrichment for LGE GABA vs TEL Glut (STAR Methods). **(J)** Representative STRING-db enriched gene sets from (I); set names paraphrased. **(K)** Two STRING enrichment networks; genes annotated with upstream TFs (**Fig S13D**). Asterisks mark genes in both sets. **(L)** CRACDL enrichment (log2FC vs. neuronal clusters from other regions; STAR Methods).

#### Regionally dispersed MGE-derived interneurons use distinct gene regulatory networks

Telencephalic interneurons derive from GE progenitors and migrate widely across the mammalian forebrain^67,68,71,72^. Consistent with their shared origin, MGE- and CGE-derived interneurons cluster by progenitor class rather than by final location at both the RNA (**Fig 1B**) and eRegulon level (**Fig 5A, Fig S13A**). MGE-derived interneurons primarily consist of *PVALB^+^* or *SST^+^*subtypes that retain shared transcriptomic identity across cortex, hippocampus, and striatum. Yet even within subtypes, they show region-associated regulatory divergence (**Fig 5B-C**).

*NKX2-1* emerged as a consistent striatum-enriched regulator for both the PVALB and SST subtypes (**Fig 5D, Fig S13B**). *NKX2-1* marks MGE progenitors and newborn neurons and is downregulated in cortex-bound interneurons^73,74^. In its TF-TF network (**Fig 5E**), *NKX2-1* activates downstream effectors (*LHX6, LHX8, ISL1, ARX, DLX1/6, PPARA*), creating a self-reinforcing circuit for striatal identity. Some effectors (*LHX6/8, ISL1*) are established *NKX2-1* targets^75,76^. Conversely, known cortical TFs (*SATB1, SATB2*, ETS-family, *MEF2C*, *TCF/NR3C1*) are enriched in cortex-localized cells and lie outside the NKX2-1 activator network^77–80^.

We also detected striatum-enriched *NFIB* in both PVALB and SST interneurons (**Fig 5D, Fig S13B**). *NFIB* activates the chemorepellents *SLIT3* and *EFNB2*, co-enriched in striatal interneuron subtypes. Slit-Robo and Ephrin signaling have previously been implicated in MGE interneuron migration^68,81^, and NFI TFs regulate GABAergic maturation^82^; these data support a role for *NFIB* in interneuron migration via Slit-Robo/Ephrin signaling.

#### Striatal MSNs and telencephalic glutamatergic neurons converge on a shared postsynaptic regulatory program

LGE-derived MSNs are striking among GABAergic neurons: in eRegulon usage, they cluster with telencephalic glutamatergic projection neurons rather than other GABAergic types (**Fig 5F-G, Fig S13A**), mirroring the RNA similarity between these two cell types (**Fig 1B**) observed in both mice and primates^20,36,37^. Despite arising from distinct progenitor domains^83,84^, MSNs and telencephalic glutamatergic neurons share eRegulons (**Fig 5H-I, Fig S13C**) including TFs implicated in both types (*EGR3, E2F3, RCOR1, FOXG1*)^85–89^, and TFs previously linked to telencephalic excitatory neurons only (*RFX3, MBNL2*)^90–93^.

Given their divergent lineage, region, and neurotransmitter identity, the LGE MSN – telencephalic glutamatergic neuron convergence points to a shared molecular function (**Fig 5H**). STRING^94^ analysis of co-enriched target genes from shared eRegulons (**Fig 5I,J**) revealed heavily overlapping, co-regulated gene sets encoding postsynaptic densities, dendritic spines, and glutamatergic synapse components with known protein-protein interactions (**Fig 5K, Fig S13D**): PSD-95/DLG-family scaffolds, SHANK-associated DLGAP/SAPAP proteins, the Ras/Rap regulator SYNGAP1, and AMPA/NMDA calcium signaling components. This shared postsynaptic program may contribute to these neuron types’ frequent co-association in disorders such as autism and Huntington’s disease^95–99^.

Beyond recovering known regulatory mechanisms, joint analysis of gene expression and regulatory elements highlights less-studied biology. *CRACDL*, which we previously identified as as enriched in striatal MSNs and telencephalic glutamatergic neurons^20^, encodes a brain-enriched, vertebrate-specific protein that localizes to the nucleus, microtubules, and centrosomes^100^. Why it is restricted to these two neuronal populations has been unclear. We identified *CRACDL* in three direct-activator eRegulons: two glutamatergic-specific (TCF4 and ZBTB18) and one (SMAD3, TGF-β pathway) also active in striatal MSNs (**Fig 5L**). In glutamatergic neurons, these TFs form an interconnected, self-reinforcing activator network upstream of *CRACDL*; only a part of this network is active in striatal MSNs (**Fig S13E**). This implicates SMAD3/TGF-β signaling in both populations, suggesting *CRACDL* is co-regulated with the spiny postsynaptic program.

### Hippocampal subfields are patterned by spatially graded gene regulatory networks

#### Spatial mapping of the marmoset hippocampal formation along its longitudinal axis

In most mammals, the hippocampal formation (HPF) lies dorsomedially below the corpus callosum – its ancestral position from cortical hem patterning^67,101^. In primates, the HPF shifts ventrally with neocortical and particularly temporal lobe expansion. We examined the marmoset hippocampus along its longitudinal (anterior-posterior, A-P) extent using our spatial data. HPF structures were detected in 25/47 MSCA spatial sections. 11 Groups and 104 Clusters mapped to the HPF, with six Groups reflecting subfield boundaries (**Fig 6A**).

**Figure 6.**
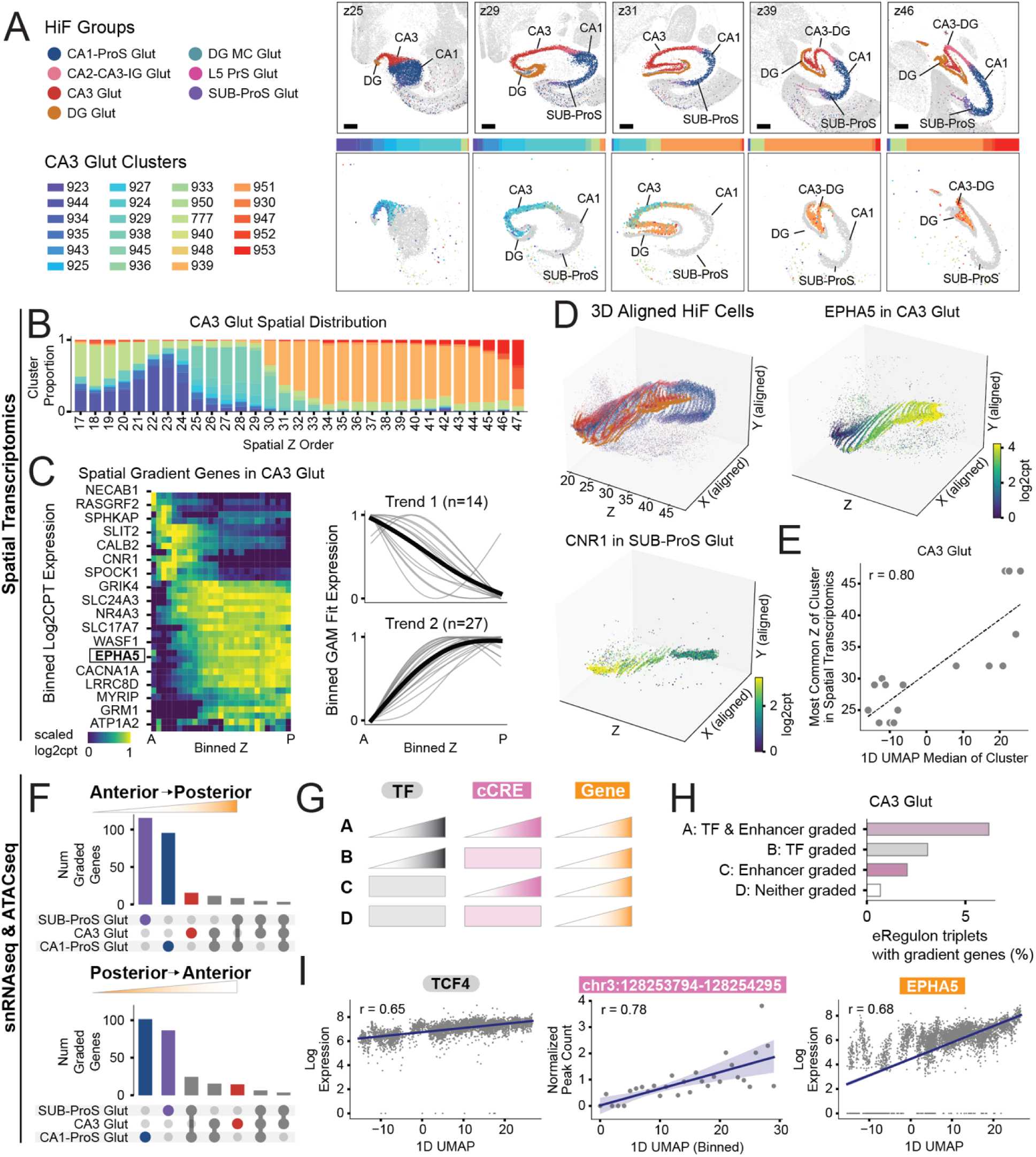
Epigenetic regulation of spatial gradients in primate hippocampal formation. **(A)** Representative spatial sections of Groups in hippocampal subfields (top) and Clusters in CA3 (bottom). **(B)** Distribution of Cluster proportions in CA3 Glut across the spatial z-axis; Cluster colors as in (A). **(C)** Left: log2cpt expression of CA3 Glut gradient genes across 30 A-P spatial bins; rows ordered by hierarchical clustered GAM fit. Right: GAM-fitted expression separated by trend 1 (n=14, decreasing posteriorly) and trend 2 (n=27, increasing posteriorly). **(D)** Three-dimensional representation of hippocampal cells colored by Group, and log2cpt of *EPHA5* (CA3 Glut) and *CNR1* (SUB-ProS Glut) in aligned spatial sections. **(E)** Correlation of median 1D UMAP coordinates with modal spatial z-section for each Cluster in CA3 Glut. **(F)** UpSet plots showing overlaps 1D-UMAP-correlated genes across hippocampal glutamatergic Groups, partitioned into A-P (top) and P-A (bottom) gene sets. **(G)** Schematic of all possible TF/enhancer gradient scenarios driving target genes. **(H)** Percentage of eRegulon triplets under each scenario in (G). **(I)** Example TCF4 +/+ eRegulon triplet in CA3 Glut: TF *TCF4* expression, enhancer peak accessibility (chr3:128253794-128254295), and target *EPHA5* expression plotted against 1D UMAP coordinates.

Cross-species mapping to the whole mouse brain atlas^1^ corroborated Group-level spatial mappings and suggested broad evolutionary conservation with some modifications. For example, the indusium griseum (IG) – a limbic structure adjacent to the corpus callosum linked to the dentate gyrus (DG) via the fasciola cinereum (FC) – remains dorsal in primates despite the hippocampus having shifted ventrally. Reports variously associate IG with CA^102^ or DG^103,104^. In the MSCA, Cluster_955 (CA2-CA3-IG Group) mapped spatially to the marmoset IG and to a restricted posterior CA3 location near the CA4 border, corresponding to mouse clusters 0400-0401 of the “025 CA2-FC-IG” supertype^1^ (**Fig S14A**). In marmoset, these cells spanned the A-P extent of the callosum and were posterior-restricted in CA3; in mouse, they occupied anterior callosum and dorsal hippocampus. This pattern is consistent with the known rodent-primate rotation of the hippocampal longitudinal axis and suggests IG is more closely related to the CA subfields than to DG.

#### Discrete cell types and continuous gene expression gradients pattern hippocampal subfields

Dorsal-ventral (D-V) gradients shape the topography of mouse hippocampus^1,105,106^. Mouse D-V corresponds to the longitudinal (P-A) hippocampal axis in primates and A-P gradients have been described in human hippocampus^107^. We observed marked spatial localization of glutamatergic neuronal Clusters along the marmoset A-P axis (**Fig 6A-B, Fig S14B**). A-P-restricted Clusters were found in DG, CA1, CA2, CA3, and Pro/Subiculum (ProS/SUB); for example separate CA3 Clusters occupied head (Clusters_943-945), head/body (Clusters_924-925), body/tail (Cluster_930) and tail (Cluster_949) positions (**Fig 6A-B**). Similarly A-P restricted Clusters were found in DG, CA1 and ProS (**Fig 6A**). DG Mossy cells (*CALB2*+/*PROX1*-) included two A-P spanning Clusters (Cluster_953, Cluster_957) and one head-restricted Cluster_956 (**Fig S14C**). Consistent with the primate rotation of the hippocampal longitudinal axis, anterior marmoset Clusters mapped to ventral mouse hippocampal types (e.g. marmoset CA3 head/anterior-body Cluster_926 mapped to mouse ventral-CA3 supertype 0076), while posterior marmoset Clusters mapped to dorsal mouse types (marmoset CA3 posterior Cluster_951 to mouse dorsal-CA3 supertype 0078 CA3 Glut_4) **(Fig S14D)**.

To characterize A-P gradients, we performed pairwise z-axis alignment using PASTE2^108^ which leverages spatial proximity and gene expression similarity without using cluster identity. Post-alignment results show that cells from the same Cluster occupy comparable spatial locations across sections, supporting continuous A-P transitions (**Fig 6D**). We then fit generalized additive models to the variable genes of each subfield Group within the spatial panel (STAR Methods; **Fig 6C, Fig S15A**). Across CA3 Glut, CA1-ProS Glut, and SUB-ProS Glut, most graded genes showed monotonic A-P increases or decreases. For example, EPHA5 (CA3 Glut) and CNR1 (SUB-ProS Glut) (**Fig 6D**), both validated by ISH in the Marmoset Gene Atlas^12,38^ (**Fig S15B**).

We leveraged the multiome data to extend beyond the 300-gene spatial panel. We embedded snRNAseq Clusters from each Group in a 1-dimensional UMAP and compared each Cluster’s median 1D coordinate to its modal Xenium *z*-section (STAR Methods). Median 1D UMAP positions strongly correlated with longitudinal *z* coordinates in all three subfield Groups, indicating that the 1D UMAP axis captures A-P position (**Fig 6E, Fig S15C-D**). We then identified genes with significant A-P gradients by correlating cell-level gene expression with 1D UMAP coordinates (STAR Methods). Across the three Groups, 244 genes were anteriorly and 115 genes posteriorly biased (|Spearman r|>0.5, p-adj<0.05). Strikingly, 318/359 (89%) graded genes were private to a single Group (**Fig 6F**). Comparison to human hippocampus A-P snRNA-seq at an anterior and posterior CA1 location^109^ showed significant overlap between human CA1 DEGs and marmoset CA1-ProS gradient genes (p-adj<0.05; **Fig S15E**), indicating conservation of subfield-specific A-P programs across primates.

#### Graded transcription factors and enhancers jointly maintain A-P expression gradients

Classical morphogen gradients (such as Bicoid in Drosophila) rely on concentration-dependent enhancer activity with the enhancer itself uniformly competent, while other patterning systems (such as spinal cord^110^) use spatially graded enhancer accessibility. Distinct enhancers are active at the rostrodorsal and caudoventral poles of the mouse hippocampal primordium^111^, but it is unknown how adult primate hippocampal expression gradients are maintained. We considered four TF-cCRE-target configurations (**Fig 6G**): (A) both TF and cCRE graded (reinforced gradient); (B) only TF graded; (C) TF uniform but cCRE accessibility graded; (D) neither graded (implying other mechanisms).

To test these, we extended the gradient analysis to cCRE accessibility across all eRegulons (cell-level correlation with 1D UMAP; STAR Methods). Using TFs-cCREs-target triplets, we classified each gradient target gene by scenario. In CA3 Glut, most gradient genes fell under scenario A (both TF and enhancer graded; **Fig 6H**), with the same pattern in other subfields (**Fig S15F**); scenario D (neither graded) accounted for only a small minority.

As a concrete example, we examined the TCF4+/+ activator eRegulon in CA3 Glut. *TCF4* itself is significantly A-P graded, increasing posteriorly. Its downstream target EPHA5 (Ephrin receptor) shows a matched posterior gradient (**Fig 6D,I**), and the TCF4-motif containing cCRE – whose accessibility correlates with EPHA5 expression – is similarly A-P graded. Thus, longitudinal gradients in excitatory hippocampal neurons are predominantly generated by coordinated grading of TFs and their cCREs, with additional mechanisms likely contributing.

### Marmoset thalamic GABAergic neurons share transcriptomic and regulatory signatures with midbrain populations

#### Cross-species mapping links thalamic GABAergic neurons to a conserved midbrain population

Aside from the thalamic reticular nucleus (RT), the rodent thalamus is almost entirely glutamatergic. In mouse, non-RT thalamic inhibitory neurons comprise only ∼6% of all thalamic neurons, most of which are found in the lateral geniculate nucleus (LGN)^7,112^. Primates harbor abundant GABAergic populations in both sensory and higher order thalamic nuclei (30-50% in some nuclei)^113,114^. From our spatial data, 20% of marmoset thalamic neurons (excluding RT, HN, dLG) are GABAergic (**Fig 7A-B**).

**Figure 7.**
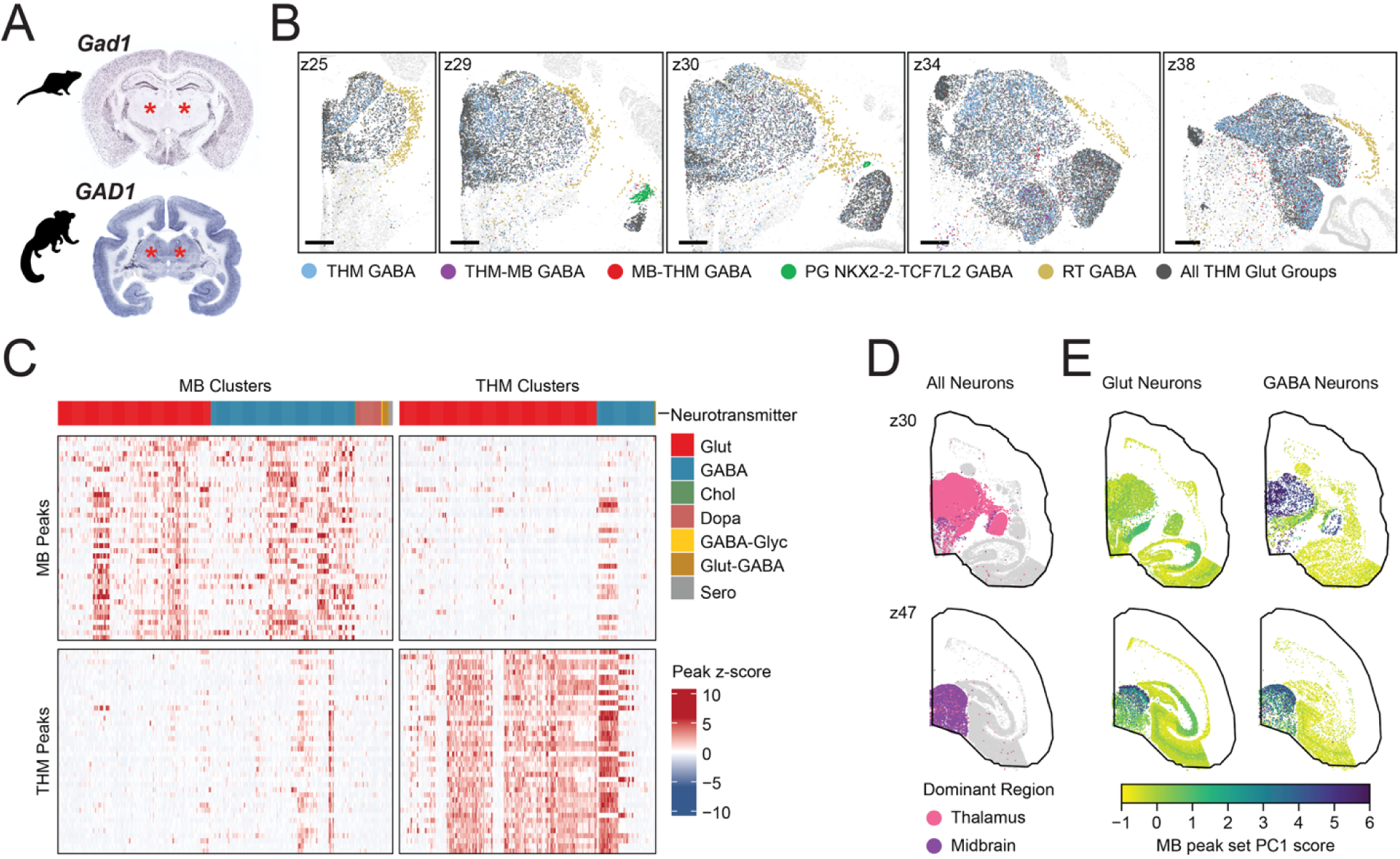
Evolutionary modifications to primate thalamus. **(A)** ISH with *Gad1* (mouse) (RRID:SCR_002978) and *GAD1* (marmoset)^12,38^ probes showing absence and presence of thalamic GABAergic cells, respectively; thalamus marked by red asterisks. **(B)** Representative spatial z-planes of neurons in the five THM GABA Groups (colored) and THM Glut Groups (grey). **(C)** Heatmap of regional marker peaks (shared in localized Glut & GABA neurons) across MB- and THM-localized clusters. **(D)** Representative spatial z-planes of all THM (top) and MB (bottom) neurons. **(E)** The same z-planes as (D), showing PC1 score of the MB peak set in glutamatergic (left) and GABAergic (right) neurons.

Prior work suggests the expanded primate thalamic GABAergic population may arise from multiple origins: diencephalon and telencephalon^115,116^, midbrain and diencephalon^7^, or midbrain, diencephalic and telencephalic (in humans)^117^. Because adult transcriptomes strongly retain lineage information^1,3,20^, transcriptional similarity between thalamic GABAergic neurons and other cell types may indicate their origins.

Non-RT thalamic GABAergic cells comprise three Groups: THM GABA, THM-MB GABA, and MB-THM GABA (**Fig 7B**). Some Clusters were nucleus-specific (e.g. LGN-restricted Cluster_38; **Fig S16A**), but many were distributed across both thalamus and midbrain (**Fig S16B**): 26% of cells in these Groups were found outside the thalamus. This thalamic-midbrain sharing raised the possibility that primate-expanded thalamic GABAergic populations derive from a conserved midbrain population. Supporting this, all three Groups aligned to a single mouse^1^ subclass, 213 SCsg Gabrr2 Gaba (**Table S5;** STAR Methods), which predominates in mouse SC with sparse LGN extension (**Fig S16C**). We validated this homology using SAMap^118^ (**Fig S16D**).

We also examined the direct mapping between the human^3^ and marmoset, plus an additional Slide-seq mouse atlas^2^. All three marmoset THM GABA Groups mapped to the human *Midbrain-derived inhibitory* Supercluster^3^ (containing samples from thalamus, midbrain, pons; **Fig 2A**, **Table S5**), and both mouse atlases mapped them predominantly to midbrain and tegmental population, reinforcing a conserved midbrain-like transcriptional identity.

#### Shared cis-regulatory elements support a midbrain origin for marmoset thalamic inhibitory neurons

We next asked whether THM GABA midbrain-like identity is also reflected at the regulatory level. By eRegulon similarity, midbrain GABAergic neurons were most similar to thalamic GABAergic neurons (**Fig S16E, S13A**). Co-enriched eRegulons included those with TFs *GATA3*, *OTX2*, *BACH1*, and *ESRRG* (**Fig S16F**).

We further hypothesized that the chromatin landscape might retain evidence of cell lineage - if some accessibility patterns are carried from an earlier developmental state. As noted earlier, midbrain glutamatergic and GABAergic neurons co-enrich for midbrain-specific peaks (**Fig S7E**). If primate-expanded thalamic GABAergic neurons originate from midbrain, they should retain some of these peaks. We ranked ATAC peaks by their shared specificity to midbrain glutamatergic and GABAergic cell types, defining midbrain-marker peaks (STAR Methods). This procedure produced distinct, regionally-specific peak sets for most brain regions (**Fig S16G**), yet many midbrain-specific peaks were also accessible in thalamic GABAergic neurons (**Fig 7E**). Projecting PC1 of this peak set onto the spatial data shows these peaks shared among midbrain glutamatergic/GABAergic and thalamic GABAergic neurons and absent elsewhere (**Fig 7F**).

Mouse developmental data suggest that excitatory and inhibitory SC neurons arise from shared multipotent progenitors^119^. In humans, *OTX2*, a shared marker of the non-RT thalamic GABAergic and SC populations, is also expressed in the midbrain and dorsal diencephalon (future thalamus) as early as gestational week 10^120^. Together with the cross-species transcriptomic and regulatory concordance, this supports a midbrain origin for most marmoset non-RT thalamic GABAergic neurons^7^, with their expansion a primate-specific adaptation.

### Interactive multimodal exploration with Cytosplore Viewer

To enable integrated exploration of the MSCA, we developed the Cytosplore ATAC Viewer as part of the Cytosplore Viewer. Complementing web-based taxonomy viewers (e.g. cirrocumulus [RRID:SCR_021646]), Cytosplore Viewer provides an offline, fast, interactive plugin-based platform for exploration of all three data modalities in depth. The viewer leverages the common taxonomy across modalities to transfer gene expression or chromatin features through common annotations, with the spatial data as a spatial reference frame. In addition, we developed a bi-directional search engine between spatial RNA-seq and spatial ATAC peak accessibility patterns (STAR Methods). Starting from a seed gene’s expression pattern across the spatial atlas, a search can be performed for ATAC peaks that exhibit a similar spatial pattern. Conversely, a spatial ATAC profile can be used as a seed to search for genes exhibiting a similar spatial expression pattern. These searches can be confined to manually selected spatial regions of interest. Cytosplore ATAC Viewer enables interactive PCA computation from user-defined sets of ATAC peaks or genes, followed by spatial visualization of PC1 (**Supplemental Movie,** reproducing **Fig 7E**). Cytosplore Viewer can be downloaded from https://viewer.cytosplore.org and is an integral part of the BICAN data tooling ecosystem.

### Discussion

#### A spatially resolved multimodal atlas of the primate subcortex

Brain cell atlases now describe cellular diversity at high resolution, but variable sampling strategies across studies mean gaps remain. In the present study, the combination of anatomically comprehensive tiling, spatial localization, and paired chromatin accessibility in a primate creates a distinct resource that is well suited for linking molecular taxonomy to anatomical context and candidate regulatory architecture. The analyses presented here highlight several ways this framework can be used to understand broad principles of molecular organization, but are only a snapshot of what the MSCA supports given the breadth of the taxonomy and its spatial and chromatin modalities.

A notable outcome is that Groups and even Clusters of the snRNA-seq defined taxonomy mapped to anatomically precise locations. This suggests that non-optimized gene panels with modest numbers of genes can still capture substantial cellular diversity. More broadly, defining cell types through quantitative variation across many genes — rather than through highly specific individual markers — aligns with the concept of cell types as peaks in a joint expression probability distribution^121^, a framework that may be especially powerful for cross-species comparisons of divergent cell populations^4,122^.

#### Distinct regulatory architectures can produce similar transcriptional results

By comprehensively surveying broad anatomical structures, our multimodal atlas clarifies not only where transcriptionally related cell types occur, but how those identities are implemented through gene regulatory programs. Although chromatin accessibility and gene expression are broadly aligned across the taxonomy (**Fig 1D**), their occasional discordance is informative: similar transcriptional outputs can arise through distinct regulatory architectures, as illustrated by shared expression of *MOXD1* in neurons and astrocytes. This case shows that transcriptional similarity cannot be assumed to predict enhancer activity across cell types – a consideration for the design of cell-type-specific genetic tools.

#### Regulatory models reveal how spatial identity is preserved or reshaped

Because postmitotic brain cells usually remain within their developmental compartment of origin, regional identity and cell type identity are typically intertwined^123,124^. But cellular identity can also be reshaped: by migration (as in MGE-derived interneurons) or by cell-type-specific functional requirements (as in telencephalic glutamatergic neurons and MSNs). Our analyses nominate TFs that may act as master regulators of these shifts, though adult data alone cannot resolve the underlying developmental mechanisms.

Our analysis of the hippocampus extends this framework to continuous regulatory variation within an anatomically defined population: discrete subfield-restricted cell types are superimposed on anterior-posterior programs marked by graded transcription factors and coordinately graded cCREs. Whether these subfield-specific gradients are coordinated by a shared upstream regulator, and how they are established during development, remain open questions.

#### Adult regulatory states generate testable hypotheses about developmental origin

Differential migration of conserved cell types is a potent route by which evolution reorganizes brain circuits^5,18^; our atlas enables predictions about developmental origin from adult regulatory state, where migration or anatomical redistribution obscures lineage relationships in single-modality analyses. The primate-expanded non-RT thalamic GABAergic population illustrates this, aligning with conserved midbrain populations across transcriptomic mapping, eRegulon usage, and chromatin accessibility. Developmental validation will be required to confirm the proposed migratory origin; future work will clarify how and when such migration occurs and its functional consequences. More generally, similar peak- and gene-set-based analyses can interrogate candidate lineage relationships, regionally patterned cell states, or disease-relevant regulatory features across the atlas. Users can project their own feature sets in our Cytosplore Viewer for interactive exploration.

#### Limitations of the Study

While comprehensive in the subcortex, the present study is anatomically incomplete. Coverage of the cerebellum and neocortex will be required for a complete taxonomy. Our limited donor number and the absence of developmental time points preclude analysis of inter-individual variation, sex differences, or developmental trajectories between cell types. Future whole-brain and developmental primate atlases, including those from BICAN, will extend this work.

Furthermore, GRN inference remains technically limited. A limitation of SCENIC+ is its insensitivity to repressive relationships^54^; accordingly, we recovered relatively few repressors (STAR Methods). SCENIC+ also relies on conventional motif matching for TFBS prediction; these may be improved by emerging deep-learning approaches^125,126^.

## Conclusion

The MSCA links high-resolution molecular taxonomies to three-dimensional anatomy across the primate subcortex. Through cross-species embedding, we disambiguate heterogeneous human cell populations and refine spatial annotations in complex anatomical subdivisions. The concurrent creation of a cross-species basal ganglia taxonomy (marmoset, macaque, and human)^32,33^ using parallel nomenclature and annotation suggests that much of the MSCA will prove conserved across primates. Ongoing work in BICAN to define cell types across the whole brain in humans and non-human primates will benefit from MSCA annotations as data collection and processing proceeds across other species. A powerful future application will be to map human genetic risk variants for neuropsychiatric and neurodegenerative disease onto spatially resolved primate cell types^127^, leveraging the spatial context to interpret noncoding variation. As such, this cellular-resolution atlas of the compact marmoset subcortex is well situated to bridge our understanding of rodent and human cell-type organization.

## STAR Methods

Resource availability

## Lead contact

Fenna Krienen

## Materials availability

This study did not generate new unique reagents.

## Data and Code Availability

Raw 10X Multiome data can be accessed at NeMO (RRID:SCR_016152, https://assets.nemoarchive.org/col-vdk0cmw) and all processed data will be publicly accessible (https://brain-map.org/consortia/hmba, https://allen-hmba-releases.s3.us-west-2.amazonaws.com/index.html#marmoset_subcortex/). Code used in the analyses in the manuscript can be found at https://github.com/KrienenLab/MSCA.

## Experimental model and subject details

### Animals

Marmoset experiments were approved, overseen, and conducted in compliance with the Massachusetts Institute of Technology IACUC under protocol number 2303000479.

## Method details

### Tissue processing

Adult marmosets (5 individuals, 3 male and 2 female, 2-7.5 years old with a median age of 5.6 years; **Table S1**) were initially sedated with Alfaxalone (12mg/kg, 10 mg/ml) and Midazolam (0.3 mg/kg, 5mg/ml) via intramuscular injection. They were further sedated with a secondary dose of Alfaxalone (4mg/kg, 10mg/ml) to ensure deep sedation followed by an intravenous injection of Euthasol (>120mg/kg, 390 mg/ml). When respiration and the pedal withdrawal reflex were eliminated, the marmosets were transcardially perfused with ice-cold carbogenated *N*-methyl-D-glucamine (NMDG) artificial cerebrospinal fluid (aCSF) (92 mM NMDG, 2.5 mM KCl, 1.25 mM NaHßPOß, 30 mM NaHCOß, 20 mM HEPES, 25 mM glucose, 2 mM thiourea, 5 mM sodium L-ascorbate, 3 mM sodium pyruvate, 0.5 mM CaClß·2HßO, and 10 mM MgSOß·7HßO; pH 7.3-7.4 adjusted with HCl). Brains were extracted into ice-cold NMDG aCSF. The brainstem, cerebellum, and olfactory bulb were manually separated using a sterile dissecting knife. The brainstem was bisected sagittally and then cut into 3-4 mm sections at an angle perpendicular to the long axis. The cerebellum was bisected sagittally and then cut into 3-4mm sagittal sections, ensuring to leave the deep cerebellar nuclei, vermis, and cerebellar hemispheres intact.

The remaining cerebrum was placed dorsal-side-down into a custom-made steel marmoset brain matrix for sectioning in ice-cold HEPES-sucrose solution (110 mM NaCl, 10 mM HEPES, 25 mM glucose, 75 mM sucrose, 7.5 mM MgClß·6HßO, 2.5 mM KCl; pH 7.4 adjusted with NaOH; ∼350 mOsm). Coronal sections were made using a razor blade at 2mm intervals from posterior to anterior for multiomic samples and 5mm intervals from posterior to anterior for spatial transcriptomic samples. The slabs were arranged in order on a teflon coated metal plate, ensuring that the posterior surface of each slab remained face-up. The metal plate with the brain samples was submerged into a shallow tray of isopentane chilled on dry ice, freezing the samples in less than a minute. Samples that became submerged were briefly air-dried on the chilled plate before further handling. From this point, all tissue was maintained and transported on the metal plate in contact with dry ice. Smaller samples (e.g., blood, olfactory bulb, dura, meninges, choroid plexus) were frozen by floating 1.5 mL tubes in the chilled isopentane. Each brain slab was vacuum sealed on dry ice and then stored at -80°C.

Subcortical dissections from each slab was performed by outlining prospective tile boundaries on each frozen slab. We intentionally sampled the following subcortical structures: the non-isocortical telencephalon (HiF, OlfC, CNU), diencephalon (THM, HY), midbrain, and majority of the hindbrain (excluding cerebellum). One to eight contiguous tiles (14-60 mg each) were dissected from each slab. Microdissected tiles were stored in microcentrifuge tubes at -80°C or used immediately for nuclei isolation. When possible, tile boundaries respected visible anatomical boundaries on the slab face. However, each tile invariably encloses multiple anatomical structures and some structures from outside the regions above are inadvertently sampled. Tiles were generally sampled from the left hemisphere of each slab; the right hemisphere was reserved for resampling due to sample or library preparation failures.

### Nuclei isolation

Single nuclei isolation of marmoset samples for 10x sequencing was conducted as described in (dx.doi.org/10.17504/protocols.io.5qpvok1exl4o/v2). Briefly, frozen marmoset neural tissue microdissections ranging from 14 to 60 mg were placed into a 3 mL dounce homogenizer containing 2 mL homogenization buffer (250 mM sucrose, 25 mM KCI, 10 mM Tris pH 8.0, 5 mM MgCl_2_, 0.1% Triton X-100, 1X Protease Inhibitor Cocktail (Halt™), Protector RNase Inhibitor, and 0.1 mM DL-Dithiothreitol (DTT)). Tissue was homogenized with approximately 10 to 20 strokes of the pestle and passed through two 15 mL conical tubes with 70 and 30 μm cell strainers. To pellet the nuclei, the sample was centrifuged at 900 x g for 10 minutes at 4°C. Nuclei were resuspended in 3% blocking buffer (1X phosphate-buffered saline (PBS), 30% bovine serum albumin (BSA), and Protector RNase inhibitor). To remove myelin from the sample, an equivalent volume of 42% OptiPrep™ density gradient medium (iodixanol) solution was added to the sample. This resulting 21% iodixanol mixture was layered on top of an equivalent amount of 25% iodixanol solution. Samples were centrifuged at 8,000 x g for 15 minutes at 4°C followed by removal of the myelin at the surface and the remaining supernatant. Samples were resuspended in 1 mL of blocking buffer and nuclei were incubated in mouse anti-NeuN PE and rabbit anti-Olig2 Alexa Fluor 488 for 30 minutes. After staining, the samples were spun down at 400 x g for 5 minutes at 4°C. The supernatant was carefully removed so as to not disturb the pellet and resuspended with 300 μL blocking buffer and DAPI (4′,6-diamidino-2-phenylindole) at a final concentration of 0.1 μg/mL.

### Fluorescence-activated nuclei sorting (FANS)

Nuclei suspensions were sorted with the Sony MA900 cell sorter using a 130 μm sorting chip. The gating strategy included first gating on their size and scatter properties. Nuclei were then gated based on DAPI signal to label DNA-containing nuclei and exclude debris and doublets. Finally, nuclei were sorted three ways based on the NeuN positive, Olig2 positive, and double negative signals. It took approximately 25 minutes to sort through each 300 μL nuclei suspension. Typically, Olig2 positive nuclei were the most abundant population, followed by slightly fewer NeuN positives, and a much smaller double negative count.

### Sample preparation for multiome sequencing

In order to enrich the neuronal nuclei populations for sequencing, the sorted nuclei were pooled at an ideal ratio of 70% NeuN positive, 10% Olig2 positive, and 20% double negative. If the resulting pool contained less than 40,000 total nuclei, the proportions of each population were manipulated to allow for a greater sample size while attempting to maximize NeuN positive input. This usually meant decreasing the proportion of double negative nuclei in the pool. After pooling, BSA was added to a final concentration of 3% followed by centrifugation at 600 x g for 10 minutes at 4°C. The supernatant was decanted using a P1000 pipette leaving approximately 250 μL above the invisible pellet of nuclei. An equal amount of 2X nuclei buffer (10X Genomics PN2000153; with Protector RNase inhibitor, 0.2 mM DTT) was added to the sample. The sample was finally centrifuged at 600 x g for 5 minutes at 4°C. The supernatant was again decanted carefully leaving just 10 μL at the bottom per desired 10X Multiome reaction. Nuclei were quantified using a hemocytometer on an EVOS M5000 fluorescent microscope. A sample concentration of at least 3,230 nuclei per μL was ideal for the 10X Chromium for a recovery of 10,000 cells/μl.

GEM generation and library preparation were performed following the manufacturer’s protocol (10x Chromium Next GEM Single Cell Multiome ATAC + Gene Expression Reagent Kits user guide, CG000338). Libraries were validated and normalized for pooling using the 4150 TapeStation System (Agilent, #G2992AA) and the Qubit Flex Flourometer (Thermo Fisher, #Q33327). Libraries were shipped to the Broad Institute and sequenced using a NovaSeq X Plus 25B flow cell (Ilumina) at a depth of 120,000 reads/nucleus.

### Xenium spatial transcriptomics data collection

Experimental details for this dataset are also described in Hewitt, Turner, et al.^33^ A fresh whole brain from a 5-year-old male marmoset (**Table S1**) had its brainstem and cerebellum resected at an oblique angle, and the remaining hemispheres were slabbed into six, 5-mm thick, coronal slabs from rostral-to-caudal, which were then flash frozen. Slabs were then divided into hemispheres, and slab face images were used to identify the three right hemisphere slabs (slab 42, slab 43, and slab 44) that contained the rostral-caudal extent of all basal ganglia structures. Specifically, the rostral pole and caudal-most portion of the caudate (approximately interaural +13.80 mm and +0.80 mm, respectively^128^) were used as anatomical landmarks for sectioning. Selected slabs were sectioned on a Leica cryostat at 10 µm onto 10X Xenium slides and collected at 200 µm intervals. During sectioning, tissue sectioned from the front of each slab was discarded until a near-complete hemisphere was obtained; similarly, incomplete sections at the end of each slab were discarded. 22 sections were collected from slab 42, 21 sections from slab 43, and 16 sections from slab 44 for a total of 59 sections. 12 sections were discarded due to Xenium instrument failure or per-section QC failure. These factors led to a final number of 47 sections (19, 14, and 14 sections per slab, respectively), which span the extent of the basal ganglia with a finest inter-section interval of 200 µm, some inter-section intervals of 400-600 µm, and larger intervals between slabs. Spatial transcriptomic sampling ended rostral to the caudal-most multiome slabs and did not include the medulla, pons, or cerebellum.

The 300 gene panel was constructed from a combination of computationally derived and manually selected marker genes. Literature derived marker genes, including genes chosen for variation between caudate and putamen^12,38^, along with shared genes from macaque and human spatial gene panels, formed a starting gene list. This list was expanded using mFISHtools^129^ and GeneBasis^130^ applied to variable genes in BG and thalamus cells from a previous marmoset census dataset^20^. Final gene panel selection was made iteratively with 10X to achieve compatibility with Xenium probe and detection constraints. Sections were fixed, probe hybridized, amplified, and imaged according to standard 10X Xenium protocols. The 10X Xenium Cell Segmentation Kit was used; the on-instrument cell segmentation algorithm was run with default settings.

Quantification and statistical analysis

### Marmoset genome reference

All sequenced libraries were aligned to “mCalJa1.2.pat.X_mitos2”^131^, a version of NCBI GenBank/RefSeq GCF_011100555.1/GCA_011100555.2 modified to contain mitochondrial gene annotations generated using the mitos2 webserver^132^ and mitochondrial chromosome CM021961.1^133^.

### Multiome sequence alignment and library QC

10x Multiome libraries were aligned using 10X cellranger-arc (v2.02). For library QC, we used cellranger-arc output metrics and MultiQC^134^ (v1.30). Out of 159 total alignments, we considered 12 RNA libraries to be QC fails for either failing to align (n=1), having <50% of reads map confidently to transcriptome (n=3), <60% of reads in cells (n=4), or >55% of reads contain the TSO sequence (n=8). We additionally failed one more library after determining a large fraction of its cells were found in low quality (“junk”) clusters. We considered 15 ATAC libraries to be QC fails for having >10% of read pairs unmapped (n=1), <50% of high-quality fragments in cells (n=7), or having <20% of transposition events in peaks be in cells (n=7). In total, 133/159 libraries passed all QC, however we maintained an additional 13 RNA libraries whose ATAC counterparts were called QC failures.

### Multiome RNA processing pipeline

RNA count matrices from cellranger-arc were processed using cellbender^135^ (v0.3.0) to remove ambient RNA. scDblFinder^136^ (v1.21.0) was used to annotate doublets using an expected doublet rate of 0.01 per 1000 cells (after removing cells with <200 UMIs). After doublet removal, we kept cells if they had 2000-50000 UMIs, 1000-13000 genes, and <3% mitochondrial gene content. Basic cell curation was done in Seurat^137^ (v5.0.3) before exporting to h5ad.

Scanpy^138^ (v1.10.4) was used for the calculation of top 4000 highly variable genes (HVGs) with “seurat_v3” as flavor and donor name as batch key. Using the HVG subset, we corrected for donor batch effects by training a scVI^34^ (v1.2.2.post2) model with donor name as the batch key and n_hidden=256, n_latent=64, n_layers=3 as the network parameters for 300 epochs to integrate cells across all donors. In the donor-integrated scVI latent space, cells were clustered using the transcriptomic clustering package^35^ (v1.0.0) with the same clustering parameters as described in the companion cross-species BG taxonomy paper^32^. Initial iterative clustering yielded 1811 clusters and additional manual curation was performed to remove 74 low quality clusters with high mean doublet scores. For each cell, raw gene expression counts were normalized with a target sum of 1e6.

Pseudobulked “metacells” for each Cluster and Group were generated by summing raw reads, which were then CPM-normalized. Normalized metacells for all Groups were reduced to only include HVGs as input for hierarchical clustering using FactoMineR::HCPC^139^ (v2.11). The resulting dendrogram was split into neurons and nonneurons, and visualized using ggtree^140^ (v3.6.2) and ggnewscale^141^ (v0.4.10). The Cluster-level metacells are used throughout other analyses. When gene expression is shown with z-scores in heatmaps, the z-scores are calculated across all Clusters, and when shown for some grouping of Clusters (for instance, for a Class, Group, Dominant Region, etc.), the Cluster-level z-scores are averaged (arithmetic mean).

### Marker gene identification and Differential Gene Expression Analysis

Marker genes for every Cluster and Group were computed using NSForest^142^ (v4.1) with default parameters. When directly comparing between 2 clusters, we used scanpy.tl.rank_genes_groups with wilcoxon rank sum tests. When comparing groupings of multiple Clusters, differentially expressed genes were calculated using DESeq2^143^ (v1.46.0) using the individual Cluster metacells (pseudobulks) as pseudo-replicates. For DESeq2, pseudobulk count matrices were subset to contain only the clusters being compared. Where noted, shrunken log2FoldChange estimates were produced using apeglm^144^ (v1.28.0).

### Cross-species mapping against reference dataset

To generate the taxonomy, we mapped our RNA dataset to various published atlas-level scRNA-seq and snRNA-seq datasets in human, mouse and other primates. MSCA dataset was restricted to 1:1 orthologous genes in the reference species. For large reference datasets with over 1 million cells, we used MapMyCells^39^ (RRID:SCR_024672) to predict labels from reference datasets to each cell in MSCA due to memory constraints. All other reference datasets were integrated with corresponding MSCA subsets using scVI with custom tuned network parameters and donor labels as covariate. KNN classifiers were then used to predict labels for each cell in MSCA based on the integrated latent space. Detailed algorithms and parameters used for each reference dataset and the associated mapping results from each reference dataset are summarized per Cluster and Group in **Table S4**-**5**.

Additional pairwise mapping among whole brain human^3^, mouse scRNA-seq^1^ and mouse Slide-seq^2^ atlases was performed using MapMyCells to identify homologous clusters across species.

### Xenium spatial transcriptomics processing and QC

Data processing and quality control for the Xenium spatial transcriptomics data were done as described in Hewitt and Turner et al^33^. Briefly, on-instrument cell segmentation was run with default settings. Segmented cells were deemed low quality and removed if they had fewer than 20 total transcripts and fewer than three unique genes (**Fig S1G**). Gene counts were normalized to log2 counts per thousands (log2cpt).

### Cross-modality cell-type mapping between spatial transcriptomics and multiome RNA-seq

Cells from the Xenium spatial transcriptomics datasets were mapped to the snRNA-seq “v4 Clusters” using the MapMyCells^39^ (RRID:SCR_024672) flat mapping algorithm with bootstrapping (100 iterations, bootstrap factor 0.95) and using all 300 genes from the gene panel (**Table S2**) as marker genes. The higher levels of the MSCA taxonomy (Class, Group) were assigned to each Xenium cell using label transfer via its mapped Cluster identity.

### In silico validation of gene panel performance for spatial RNA-seq mapping

MapMyCells was used to self-map the multiome RNA to their own taxonomy using the same settings as was used for the Xenium spatial RNA-seq mapping. The gene sets used for the mapping task were: original 300 gene spatial panel; the 4000 HVGs used for multiome RNA-seq scVI integration and clustering; Group- and Cluster-level NSForest markers from the taxonomy (n=398 and n=2916, respectively); all marmoset transcription factors, taken from AnimalTFDB v4.0^145^ (n=1329); the mouse ABC atlas MERFISH panel genes (n=500); and all genes used for labelling in the mouse ABC WMB atlas taxonomy (n=194) (**Table S2**). For assessing performance, the original cell labels in the multiome taxonomy were used as ground truth for calculating precision, recall, and F1 scores.

For the spatial gene panel feature importance analysis, a Random Forest classifier (sklearn v1.8.0) was trained on the multiome snRNA-seq data using the union set of the 4000 HVGs and the 300 Gene Spatial Panel (n=4104). The model was trained with 1000 decision trees, and impurity-based feature importance scores were extracted to quantify the extent to which each gene contributed to splitting decisions across the forest. Training was parallelized in 40 batches of 25 trees each.

### Generation of MSCA Taxonomy

Groups in the initial draft taxonomy were generated by aggregating clusters with similar transcriptomic signature and consistent mapping to the same reference label in all relevant reference datasets. After mapping snRNAseq defined Clusters to Xenium spatial transcriptomics data as described above, we examined each Cluster’s spatial distributions and refined Cluster-to-Group assignments. We merged Clusters with similar transcriptomic profiles and contiguous spatial localization, as well as split transcriptomically similar Groups into subsets with clearly distinct spatial territories. All acronyms used in Group names are provided in the Harmonized Ontology of Mammalian Brain Anatomy (HOMBA, RRID:SCR_027628)^146^. Groups were then assigned to the corresponding Class derived from the mouse taxonomy^1^. Marker genes that are consistently expressed in MSCA and reference datasets are included as part of the Group name. Clusters identified to be in the basal ganglia were assigned a Group based on the cross-species consensus taxonomy generated in Johansen, Fu, Schmitz et al.^32^. In the present manuscript, two BG Groups, STRv D1 NUDAP MSN and AMY-SLEA-BNST GABA, were renamed to AMY-STR ITC-NUDAP MSN and CEN-SLEA-BST GABA, respectively, to reflect their anatomical distributions when considering locations outside of the BG.

Two Groups (OB Dopa-GABA and OB FRMD7 GABA) received olfactory bulb (OB) anatomical labels due to their shared gene expression programs with OB periglomerular interneurons. However, cells from these Groups localized to the white-matter boundary of the striatum in the spatial transcriptomics donor, and these cells are unlikely to have originated from the OB itself because we did not sample the rostral-most slab, where the OB is located, in the multiome donors. These Group labels were inherited from the BG cross species consensus taxonomy, and the characterization of these two peri-striatal Groups in marmoset, as well as macaque and human, is described in greater detail in the companion paper^32^.

Additionally, we used our spatial mapping to manually assign a Dominant Region to each neuronal Group in our taxonomy where possible. For Groups with representation across multiple anatomical structures, we refined these assignments at the Cluster level.

### Comparative analysis across mouse, marmoset and human neurons

To assess the transcriptional similarity and expression variability of cell types across mammalian species, we first identified homologous cell types. We took the cross-species mapping results of mouse^1^, marmoset MSCA, and human^3^ atlases generated above and used human clusters as the common anchor. A mouse or marmoset cluster was considered homologous to a given human cluster if >70% of its cells mapped to that human cluster. Using this criterion, 114 of 382 human clusters were identified as having homologs in both mouse and marmoset. Each dataset was then restricted to the corresponding set of 1:1 orthologous genes, and these subsets were used as input to quantify expression similarity per cluster using expressolog^51^.

### Multiome ATAC processing pipeline

ATAC processing was done following RNA analysis and the generation of a draft taxonomy, and using only libraries and cells that passed RNA QC and cell curation (for a final 632920 ATAC cells, vs. 678917 in RNA). All ATAC preprocessing was done using SnapATAC2^49^ (v2.8.0). After generating fragment files based on our RNA-curated cells, a draft version of our subcortical atlas “Group” level annotation was used for MACS3 peak calling, and peaks were merged using “merge_peaks” (half_width=250), generating >1.7 million peaks. We generated a full cell-by-peak matrix using “make_peak_matrix” (counting_strategy="insertion") as well as a pseudobulked cluster-by-peak matrix. We focused subsequent analysis on a reduced peak set that contains only peaks with non-zero signal in >50% of the cluster-level pseudobulks in at least one Group (n=462,698 peaks).

When peak accessibility is shown in heatmaps (z-scored), they are treated the same as the RNA: z-scores are calculated across all Clusters and then averaged (arithmetic mean) as needed. bigWig files were generated for our taxonomic Groups using “export_coverage” (normalization="CPM", counting_strategy=”insertion”, bin_size=1). We used “topic modeling” (with pycisTopic^50^) to group peaks into modules and to produce a low-dimensional representation of chromatin accessibility.

### Multiome ATAC-seq browser shots

The browser shot in **Figure 3D** was generated using IGV desktop^147^ (v2.16.0) and a single raw Class-level bigWig file. The browser shots in **Figure 4F** were generated in R using github.com/mdeber/browser_shot_plotter with CPM-normalized bigWig files and a binsize of 50bp. Because no bigWig track was generated for “All Nonneurons besides Astrocytes”, that track was generated by merging all other nonneuron Group CPM-normalized bigWig files and renormalizing to sum to one million.

### Gene Regulatory Network (GRN/eRegulon) calling and scoring

Due to performance constraints, SCENIC+^54^ (v1.0a2) was run separately on 50000 neuronal and 50000 non-neuronal cells sampled using geometric sketching^148^ based on our RNA scVI latent space. We performed Topic modelling with pycisTopic^50^ (v2.0a0) using default parameters, and we chose the optimal number of topics (200 for neurons, 175 for non-neurons) based on log-likelihood. Three sets of candidate enhancers were created by taking (1) top 3000 peaks per topic, (2) differentially accessible peaks on the imputed matrix at the Group level (Wilcoxon rank sum test logFC>0.5 and Benjamini–Hochberg-adjusted Pß<ß0.05) and (3) selection by topic binarization using the Otsu method. A custom database using the reduced peak set was created and motif enrichment analysis was implemented with pycistarget^149^ (v1.1). Finally, we ran the SCENIC+ workflow with default settings. SCENIC+ eRegulons are called either “direct” in the case of an exact within-species TFBS motif match, or “extended” if based on motif similarity or orthology. In neurons we produced 321 direct (307 activator + 14 repressor) and 229 extended (214+15) eRegulons; and in non-neurons 173 direct (158+15) and 126 extended (115+11) regulons. All eRegulons shown and discussed are “direct”, with activators being classified by SCENIC+ as “+/+” and repressors “-/+”.

eRegulon activity was scored at the cluster-level using RNA or ATAC pseudobulks with the R package AUCell^54^ (v1.28.0) using default parameters. When eRegulon z-scores are used, these are generated at the Cluster-level using all clusters, and unless otherwise specified, these are based on gene AUC. When eRegulon z-scores are generalized across a set of clusters, the arithmetic mean of the Cluster z-scores is used. For Group-level threshold-based analyses of OTX2 expression (e.g., defining OTX2+ Groups in **Fig 3F**), a 30 CPM cutoff was applied. eRegulon network plots were generated in R using ggraph^150^ (v2.2.2); in network visualizations, TF-gene edges are displayed only if the SCENIC+ importance score exceeds 1.5 (**Fig 3H**); for TF-TF networks centered on a focal TF, connectivity is restricted to two hops upstream or downstream (**Fig 5E**).

When multiome data (e.g. eRegulon activity, which is gene AUC z-score) is “projected onto spatial data” (as in **Fig 3E**), each spatial cell’s mapped Cluster is taken as the cell identity, and the eRegulon gene AUC z-score for that Cluster is plotted. This means all cells with the same mapped Cluster will have the same values projected.

### Cytosplore Viewer

Cytosplore Viewer is a plugin-based visual analytics system designed to support comparative analysis of multi-modal brain atlases across species. It is built on Manivault^151^, an application building toolkit for visual analytics applications for multi-modal high-dimensional datasets. By integrating coordinated visualizations of embeddings, gene expression, spatial data, and regulatory signals, the system links local compute to the underlying single-cell measurements and enables iterative exploration of complex biological datasets. Cytosplore Viewer leverages local, on-the-fly computation in combination with the fact that ATAC-seq and RNA-seq are annotated with the same, fine-grained clustering. This enables average feature profiles for ATAC peaks and gene expression to be projected on the spatial data. By approximating features with cluster averages, real-time interaction, searches and visualization for data exploration can be achieved on standard compute hardware. This allows for spatial visualization of any of the ATAC peaks, as well as of transcriptome-wide gene expression, in two linked scatterplots.

The spatial mapping module offers a search engine for genome coordinates in the ATAC data, as well as gene symbols in the RNA-seq data, enabling the spatial visualization of these single-cell features. These views are complemented with a bar chart showing the top cell types (and proportions) in which these features are predominantly expressed. In addition, the corresponding peaks and genes are projected onto the RNA UMAP. The module also supports PCA on user-defined feature sets, where PC1 is calculated and mapped onto spatial coordinates, with optional filtering to restrict visualization to selected cell types. By integrating these operations into a single environment with low-latency user interaction, the ATAC Viewer enables interactive exploration of spatial regulatory and gene expression features, enabling user-drive hypothesis generation in a seamless analytical workflow.

For a detailed description of the design considerations and process of Cytosplore Viewer, see Chapter 5 of Basu , 2026 (https://www.dropbox.com/scl/fi/10e9lw7kjdkvye9uczbzz/Soumya_PhD_Thesis_Final.pdf?rlkey=az9v4khlvo3h5sn63aanwniem&dl=0), and viewer.cytosplore.org.

### RNA/ATAC co-enrichment analysis by region and neurotransmitter

To look for the most conspicuous overlaps of specific genes (RNA) and peaks (ATAC) across neurons by region and neurotransmitter (as in **Figure S3E**), we regrouped our neuronal clusters by dominant region and neurotransmitter. Then, using the raw Cluster-level pseudobulked metacells (using all genes for RNA, or the reduced peak set for ATAC), we ran DESeq2, treating each cluster as a pseudoreplicate observation of each region+neurotransmitter pairing, and as a background we used all neuron clusters from other brain regions. We then used apeglm to shrink the log2FoldChange estimates, which reduces the magnitude of log2FoldChanges for features with higher variance, thus producing single adjusted parameter estimates for enrichment. To look for overlaps among highly-specific features, for RNA we used the top 2% of genes by shrunken log2FoldChange, and for ATAC, the top 1% of peaks. Then, for every pairwise comparison between a region+neurotransmitter (which could be between neurons that share a region, or share a neurotransmitter, or share neither), we scored co-enrichment as the ratio of observed overlaps vs. that expected by random chance. High co-enrichment indicates that the neurons being compared share a large fraction of their most-specific features. The same DESeq2/apeglm pipeline was used to compute per-gene enrichment scores for individual groupings vs. neurons from other regions; CRACDL enrichment in **Fig 5L** shows the resulting apeglm-shrunken log2FoldChange.

### Principal Component Analysis of Astrocyte Clusters

Metacells matrix with top 4000 HVGs (see ‘Multiome RNA processing pipeline’ above) for each astrocyte Cluster was used as input for the principal component analysis (sklearn.decomposition.PCA (v1.5.2), n_components=10). PC1 and PC2 captured 27.3% and 18.2% variation respectively.

### TEL Glut vs. LGE GABA functional convergence analysis

“TEL Glut” and “LGE” are defined in **Figure 5G**, which is based on regrouping neuronal Clusters by Dominant Region, Neurotransmitter, and GE origins (see main text). The “background” set includes all other neuronal groupings. All enrichment is based on z-score differences vs. that background set, with difference >1 in both TEL Glut & LGE being used to call co-enriched direct eRegulons. Co-enriched genes within those co-enriched eRegulons were also selected using the same criteria (Cluster-level z-scores are averaged for each grouping, and differences >1 are again considered enriched). That set of 279 genes was exported, and the 242 coding genes were used for enrichment analysis in string-db (v12.0, permalink: https://version-12-0.string-db.org/cgi/network?networkId=bzt5qpNsbveL). Selected network plots were downloaded from string-db. Dot plot was recreated for selected terms in ggplot2 (v4.0.2). Enrichment values reported in **Fig 5J** are the harmonic mean of observed/expected ratio and - log(FDR). UpSet plots were generated with R package ComplexHeatmap (v2.25.2). Given the number and redundancy of significant terms, terms selected for use in final plots were chosen manually following both a clustering analysis and size/clarity considerations.

Abbreviations used in STRING gene set enrichment analysis are “GO” (Gene Ontology), “CC” (Cellular Component), “MF” (Molecular Function), “BP” (Biological Process), “NMDA” (N-methyl-D-aspartate), “AMPA” (α-Amino-3-hydroxy-5-methyl-4-isoxazolepropionic acid), “CaM” (calmodulin), and “PPI” (protein-protein interactions). Abridged term names are “Reactome: NMDA unblocking” (“Unblocking of NMDA receptors, glutamate binding and activation”), “Reactome: AMPA trafficking” (“Trafficking of AMPA receptors”), “Reactome: Ras after NMDA/Ca2+” (“Ras activation upon Ca2+ influx through NMDA receptor”), “Reactome: Postsynaptic signaling” (“Neurotransmitter receptors and postsynaptic signal transmission”), “Reactome: Neurexin/neuroligin” (“Neurexins and neuroligins”), “GO MF: CaM kinase activity” (“Calmodulin-dependent protein kinase activity”), “GO BP: Calcium import” (“Calcium ion import across plasma membrane”), and “STRING: Postsynapse & PPI” (“Postsynaptic cell membrane, and Protein-protein interactions at synapses”).

### Alignment of Xenium Spatial Transcriptomics data in three-dimensions

To align consecutive Xenium slices, we used PASTE2^108^, which leverages the spatial location and gene expression of all cells simultaneously. Due to the large number of individual cells, cells are first aggregated to alignment spots with a radius of 120um. For each alignment spot, raw gene expression is computed as the sum of expression profiles from its constituent cells. The spatial location of each spot is set to the UMI-weighted centroid of these cells, and its weight is defined as the number of constituent cells. Following standard procedures, gene expression of alignment spots are log-normalized and reduced to 30 principal components. Adjacent pairs of slices are then aligned with PASTE2 with minimum overlap fraction of 0.9, Wasserstein (spatial) loss fraction of 0.1, and marginal distribution weighted by cell counts. After computing the transport map between each adjacent pair, an affine transformation is trained to fit the transport map, minimizing squared distance between the barycentric projection and affine projection of all spots in the source slice. Finally, to register all slices into a common space, adjacent affine transformations are composed such that each slice can be projected onto the first one by a single affine transformation.

### Hippocampal Gradient Analysis

Cells from hippocampal Groups CA3 Glut, CA1-ProS Glut and SUB-ProS Glut Group in the spatial transcriptomics and multiomics datasets were included for this analysis. In the Xenium spatial transcriptomics dataset, because the hippocampus was only present from sections z=21 onwards, any clusters mapped to anterior sections were excluded as low-confidence. All subsequent analyses are conducted for each Group separately.

For examining longitudinally graded genes out of the 300 genes included in the spatial transcriptomics data, we restricted our screening to only genes with log2cpt that vary more than 0.5 standard deviations across the z-axis. To reduce sensitivity to uneven sampling density across sections, cells were binned (n_bins=30) along the z-axis using quantile binning. Bins containing fewer than 10 cells were excluded from downstream analysis. For each bin, we computed the mean z coordinate and the median expression value for each gene. To cluster continuous expression trends across spatial bins, we fit a generalized additive model (pyGAM, v0.12.0) for each gene as a smooth function of z position using 6 splines, with smoothing parameter λ = 0.6 and no intercept. Fitted expressions of each gene were used for hierarchical clustering (seaborn.clustermap v0.13.2, metric=’euclidean’, method=’ward’). Across the three subfield Groups tested (CA3 Glut, CA1-ProS Glut, SUB-ProS Glut), two dominant trends were evident, with majority genes showing monotonic increases or decreases along the A-P axis.

We then used the multiomics dataset to look for transcriptome-wide longitudinally graded genes. In the multiomics dataset, additional clusters with fewer than 25 cells mapped across the remaining spatial transcriptomics sections were considered low-confidence and were excluded. One-dimensional UMAP embeddings (1D UMAP) were computed from snRNAseq data separately for each Group using scanpy.tl.umap with n_components = 1. The median 1D UMAP coordinates of each cluster were then correlated with the modal spatial *z*-order to which that cluster mapped in the Xenium spatial transcriptomics data. To identify genes with A-P gradients in the hippocampus, we computed correlations between gene expression and the 1D UMAP coordinates in the multiome dataset. P-values for all correlations were adjusted for multiple testing using Benjamini-Hochberg; features with |Spearman r| >0.5 and Benjamini-Hochberg-adjusted p < 0.05 were considered significantly graded. For cross-subfield comparisons of graded genes (**Fig 6F**), significantly graded genes from each hippocampal glutamatergic Group were partitioned by the sign of their correlation with modal spatial z-position, yielding anterior-posterior and posterior-anterior sets. Overlaps were visualized as UpSet plots.

Similarly, we identified peaks with A-P gradients by correlating the binned chromatin accessibility (n_bins=30) against the 1D UMAP coordinates due to the sparsity of the ATAC data. Significance of peaks are assessed similarly as genes above (|Spearman r| >0.5, p-adj < 0.05). To test the frequencies of each hypothesis in **Fig 6H**, we subsetted the eRegulon triplets identified above based on graded TFs and/or peaks, and then calculated the percentage of target genes that were significantly graded within each subset.

### Cross-species mapping of primate non-RT Thalamic GABAergic population

In addition to methods described in ‘Cross-species mapping against reference dataset’ section above to find homologous rodent cell types in the mouse brain atlas using MapMyCells, we also ran SAMap^118^ following the same methods and parameters described in Corrigan et al.^4^.

### Regional ATAC marker analysis

Related to **Figure 7E-F** and **Fig S7E**, we defined regional markers in chromatin accessibility to be ATAC peaks that are highly specific to a single region, but also shared among lineage-separated neurons. For this, we restricted analysis to glutamatergic and GABAergic neurons, and to regions that contained >2 clusters of each neurotransmitter (AMY, BF, CTX, EXA, HiF, HTH, MB, THM). Using our Cluster-level CPM-normalized ATAC metacell matrix, we defined region+neurotransmitter specificity as the fraction of a peak’s normalized reads that are found within that region+neurotransmitter. This naturally had the tendency to favor peaks with at least moderate accessibility. Then, to find peaks with shared specificity across glutamatergic and GABAergic clusters, we first ranked peaks by region+neurotransmitter specificity (with 1 being the least specific), and summed these ranks for the two neurotransmitters. We took the 40 highest combined ranks as a set of regional ATAC markers.

## Acknowledgements

This publication was supported by and coordinated through the Brain Initiative Cell Atlas Network (BICAN), RRID:SCR_022794. This research was supported by the National Institutes of Health award UM1MH130981. The work was additionally supported by NIH DP2MH140136 and the Klingenstein–Simons Fellowship (F.M.K.), and the Netherlands Organization for Scientific Research NWO:024.004.012 (B.L.), and the McDonnell Fellowship in Neuroscience at Princeton University (S.D., N.W.), and National Institutes of Health 5T32MH065214 (N.W.). We thank MIT DCM veterinarian staff and animal care technicians who assisted in anesthesia, post-operative care, clinical support, and animal husbandry. We thank Christina DeCoste, Gabriel Palmieri, Jailene Garcia, and the Molecular Biology Flow Cytometry Resource Facility which is partially supported by the Rutgers Cancer Institute of New Jersey NCI-CCSG P30CA072720-5921.

## Declarations of Interest

H.Z. is on the scientific advisory board of MapLight Therapeutics, Inc. The other authors declare no competing interests.

## Declaration of generative AI and AI-assisted technologies in the writing process

During the preparation of this work the authors used ChatGPT in order to correct grammatical errors and improve readability of original drafts. After using this tool/service, the authors reviewed and edited the content as needed and take full responsibility for the content of the published article.

## Notes

### Summary of Updates

Figure 4, 5, 6 revised; author affiliations updated.

