## Supplemental Figures for "Spatial patterning of transcriptional and regulatory programs in the primate subcortex"

### Supplemental Tables

**Table S1.** Donor overview

**Table S2**. Gene sets for evaluating spatial gene panel performance

**Table S3.** Self-mapping summary

**Table S4.** Cluster-level summary of quality control statistics, marker genes, and cross-species mapping results.

**Table S5.** Group-level summary of quality control statistics, marker genes, and cross-species mapping results.

**Table S6**. eRegulon triplets identified in MSCA

**Supplemental Movie**. Tutorial of Cytosplore Viewer on analysis in Fig 7E.

Supplemental Figures

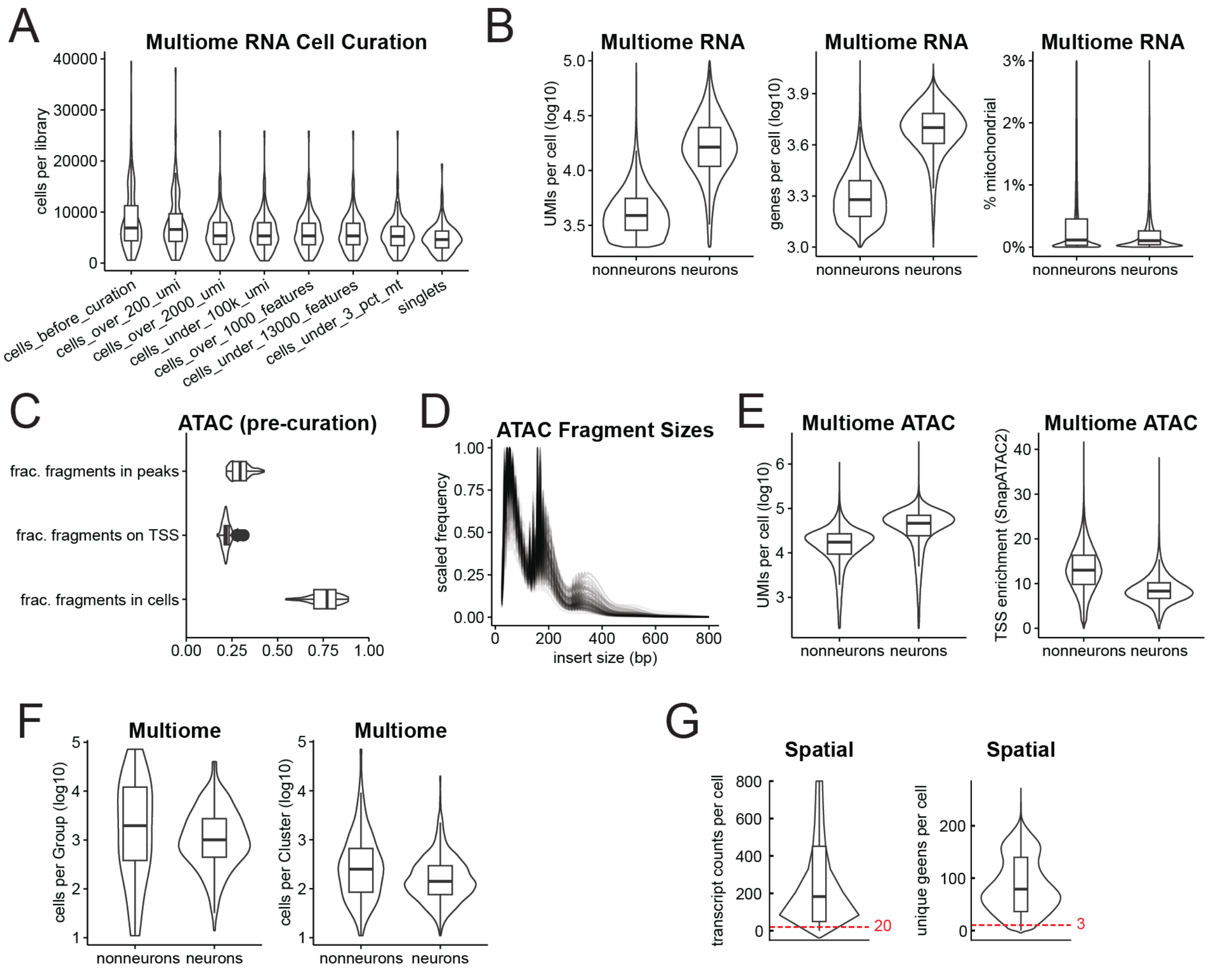


**S1. Quality metrics of multiomics and spatial transcriptomics data**, related to Fig 1. **(A)** Cell curation in multiome libraries, based on RNA metrics. Only libraries passing library-level QC are shown. **(B)** Multiome RNA metrics for curated cells. **(C)** ATAC metrics from cellranger-arc alignment summary for libraries passing library-level QC. **(D)** ATAC insert size distributions for all libraries passing QC, showing robust retention of nucleosomal fragments. **(E)** ATAC metrics for curated cells. **(F)** Multiome cells in each Group and Cluster. **(G)** Xenium spatial transcriptomics metric cutoffs for curated cells.

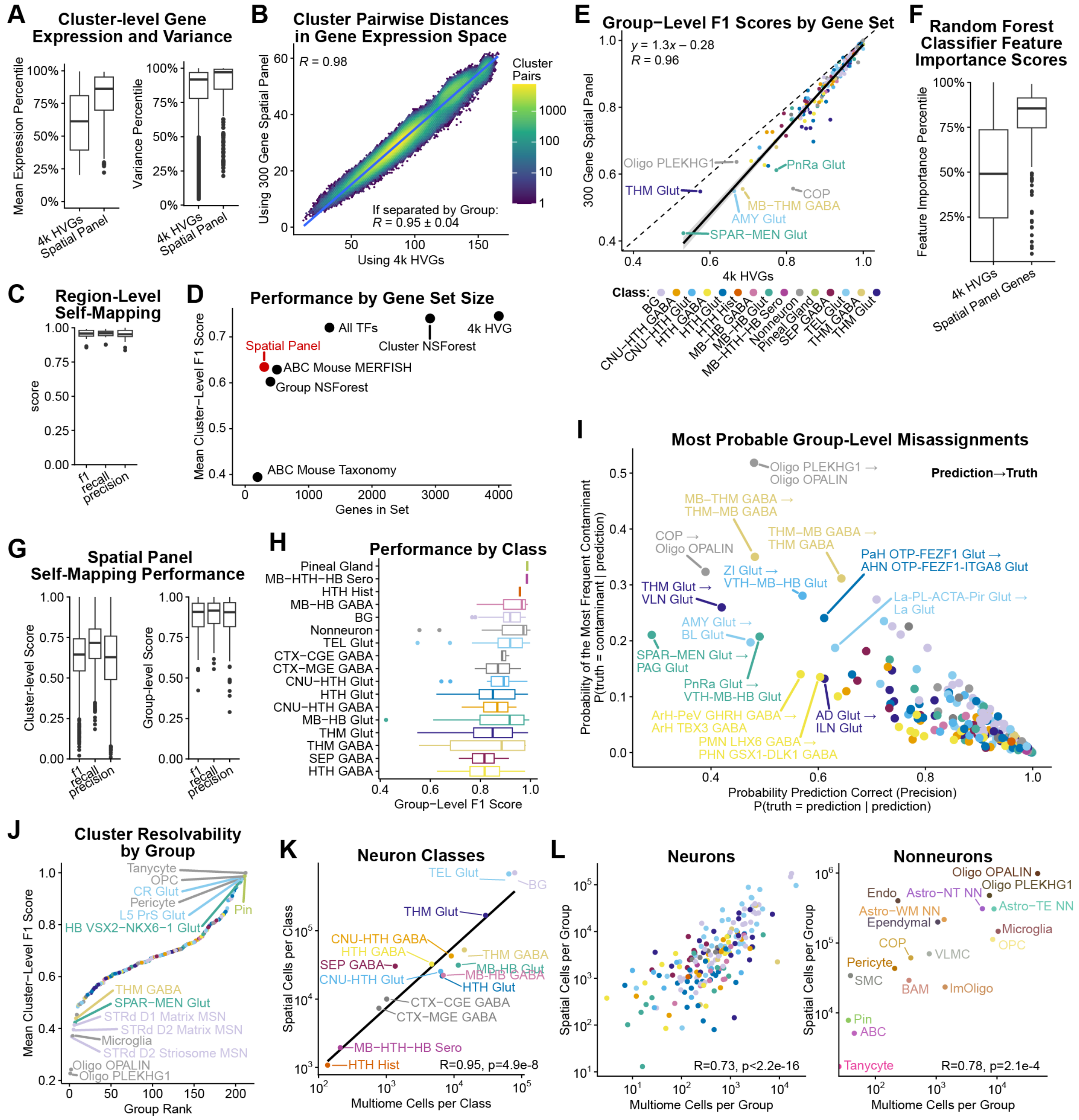

**S2. Spatial gene panel performance for localizing cell types**, related to Fig 1 and Methods
**(A)** The expression and variance of genes in the two sets is compared. Percentiles are calculated based on all genes. Spatial panel genes are both robustly expressed and highly variable across cell types. Individual clusters have 53-113 spatial panel genes within the top 1000 genes by expression. **(B)** Euclidean distances between all cluster pairs are calculated in gene expression space for the two gene sets, and those distances are plotted on each axis. High correlation indicates that the broad structure of inter-cluster relationships is

reproducible with the spatial gene panel. **(C)** Using only the 300 genes in the spatial panel to self-map the multiome RNA back to its own taxonomy using MapMyCells results in 97.5% of localized neurons receiving the same "Dominant Region" annotation. F1 score is the harmonic mean of precision and recall, which is always closer to the lower number. **(D)** The Cluster-level self-mapping performance of various gene sets are compared (**Table S2**). **(E)** Self-mapping performance at the Group-level is compared for the two gene sets. A very similar trend was observed when comparing performance at the Cluster level (y=1.2x-0.27, R=0.95). The 10 worst Groups by spatial panel mapping are labelled. **(F)** In addition to MapMyCells, a Random Forest classifier was trained on the multiome RNA and feature (gene) importance scores were extracted (see Methods). The rank-percentile is shown for each gene. **(G)** Self-mapping performance metrics for the spatial gene panel at the Cluster and Group levels. Relatively high recall in Cluster-level mapping indicates that the spatial Cluster assignments are subject to noise but mostly accurate in aggregate. **(H)** Performance at the Group-level is shown by Class, sorted by mean F1 score. **(I)** Following self-mapping with the 300 gene panel, >90% of cells receive the correct Group assignment. Highlighted here are the Groups with the lowest accuracies. For a cell that receives a given Group prediction, the probability the prediction is correct (x-axis) is plotted against the probability that the cell actually came from the most frequent contaminating Group (y-axis). For example, the uppermost point shows that 48% of cells mapped to "Oligo PLEKHG1" were actually in that Group (x-axis), while 52% were actually "Oligo OPALIN" (y-axis). Groups are colored by Class as in panel **E**. **(J)** For each Group, the Cluster-level mapping performance is averaged. Worst and best performing Groups are highlighted. In principle, some Groups' Clusters are not resolvable using the spatial gene panel and MapMyCells. Groups are colored by Class. **(K-L)** Cellular abundances are compared between the spatial donor and the most abundant multiome donor (CJ23.56.003, "Nutmeg"). Cell type abundances are largely similar for both neurons and nonneurons, although the latter was rebalanced in the multiome dataset using FANS (see Methods).

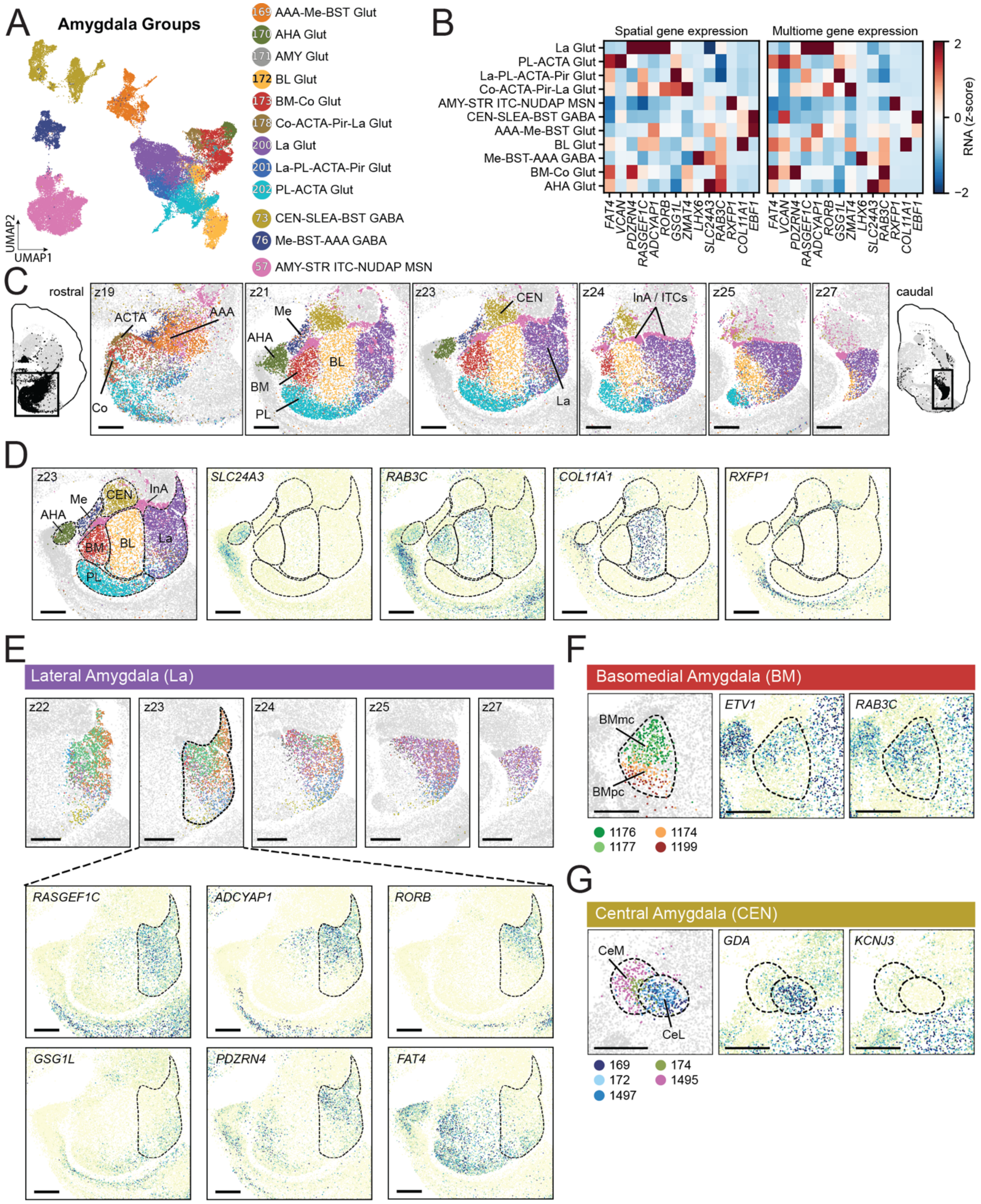


**S3. Transcriptomic Groups in the Amygdala and associated marker gene expression,** related to Fig 1. **(A)** UMAP of the twelve Groups localized to the Amygdala: nine glutamatergic Groups, two GABAergic Groups, and finally the AMY-STR ITC-NUDAP MSN Group, which localizes to both the intercalated nucleus (InA) that weaves between the other amygdala nuclei and the NUDAP compartments of the striatum. The MSCA spatial data supports the hypothesis that the intercalated cells (ITCs) that populate the InA have a shared transcriptomic–and perhaps developmental–identity with the eccentric medium spiny neurons (MSNs) found in the striatum[5]. **(B)** Marker gene expression across amygdalar Groups are consistent in spatial transcriptomics (left) and snRNAseq (right). **(C)** Representative z-planes showing spatial localization of amygdalar Groups to the major nuclei of the amygdala, including the anterior (AAA), medial (Me), paralaminar (PL), basomedial (BM), basolateral (BL), central (CEN), lateral (La), and intercalated (InA) nuclei, as well as associated transition structures, including the amygdalocortical transition area (ACTA), cortical nucleus of amygdala (Co), and amygdalohippocampal area (AHA). **(D)** Spatial distribution of representative positive regional marker genes for AHA (*SLC24A3, RAB3C*), BL (*RAB3C*, *COL11A*), BM (*RAB3C*), and InA (*RXFP1*). **(E)** Spatial localization of Clusters belonging to the La Glut Group (top), which subdivides the La nucleus along its dorsal-ventral, medial-lateral, and anterior-posterior axes. Cluster distribution is driven primarily by combinatorial overlap of complicated gene patterns, as illustrated by the complex gene gradients of *RASGEF1C*, *ADCYAP1*, *RORB*, *GSG1L*, *PDZRN4*, and *FAT4* that span the La. **(F)** Spatial localization of Clusters belonging to the BM Glut Group. The internal dorsal-to-ventral organization of clusters, which roughly maps onto reported magnocellular (BMmc) and parvocellular (BMpc) parcellations, is driven in part by differing transitions in the gradient genes *ETV1* and *RAB3C*. **(G)** Spatial localization of Clusters belonging to the CEN Glut Group. The distinction between the medial (CeM) and lateral (CeL) subdivisions of the CEN nucleus is driven in part by the combination of a step-like transition in *GDA* expression and the absence of *KCNJ3* expression in the CeL.

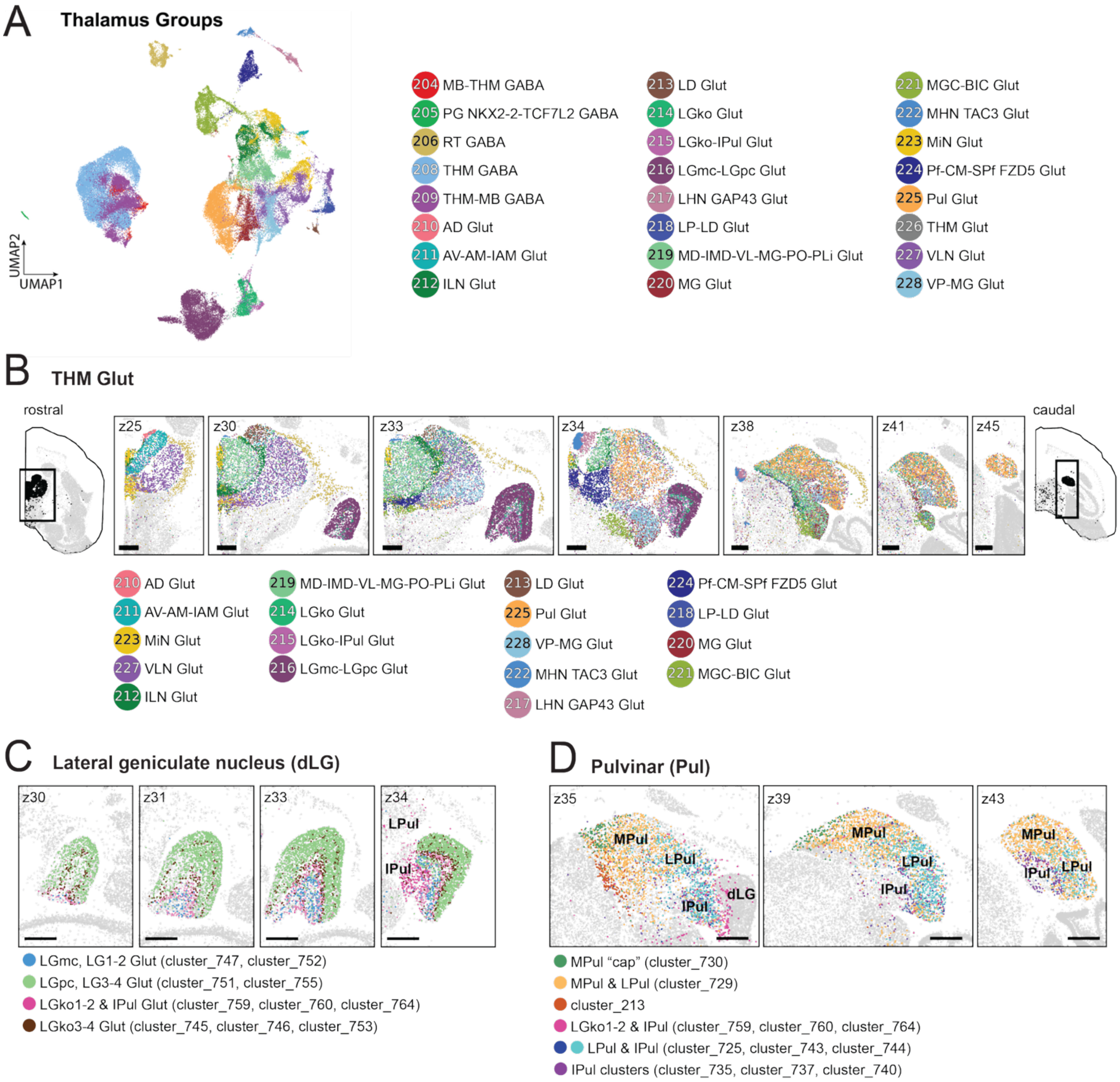


**S4. Spatially distinct transcriptomic Groups in the Thalamus**, related to Fig 1. **(A)** UMAP of all the 24 Groups in the Thalamus, including 19 glutamatergic Groups and five GABAergic Groups. **(B)** Representative spatial sections of the 19 Groups in the THM Glut Class. **(C)** Representative spatial sections showing the laminar organization of the dorsal lateral geniculate nucleus (dLG) revealed by MSCA Clusters. In primates, the dLG is a four- (marmoset) or six-layered (macaque, human) structure; whereas, in rodents, it has a core-shell organization[152,153]. LGmc (magnocellular, LG1-2): Cluster_747, Cluster_752; LGpc (parvocellular, LG3-4): Cluster_751, Cluster_755; LGko1-2 (koniocellular) & IPul shared clusters: Cluster_759, Cluster_760, Cluster_764; LGko3-4: Cluster_745, Cluster_746, Cluster_764. **(D)** Representative spatial sections of the Pulvinar nucleus, which is greatly expanded in primates. Previous work indicates that by the perinatal period, nuclei of the marmoset pulvinar exhibit gene expression differences not found in mouse pulvinar[12]. Unlike the inferior and lateral pulvinar (IPul, LPul), the medial pulvinar (MPul) – a specialized nucleus of the primate thalamus that connects to prefrontal cortex and may be involved in distinct aspects of visual function – does not have a clear anatomical homologue in rodents[10,154]. Consistent with molecular distinctions across marmoset pulvinar nuclei, Cluster_729 is biased to MPul, while Cluster_728 is biased to LPul. Direct comparison of Cluster_728 and Cluster_729 revealed differential expression of 486 genes with more than two fold enrichment in either Cluster ($p$-adj $< 0.05$). Notably, among them are genes reported previously to delineate medial and lateral pulvinar, such as *CALB1* and *NNAT*[12,155,156]. While there are clusters specific to IPul (Cluster_735, 737, 740), many clusters appear shared across LPul and IPul (Cluster_725, 743, 744), perhaps hinting at a gradient-like organization to the pulvinar. Two clusters notably localized to specific regions within MPul: Cluster_730 to a dorsal-medial "cap" and Cluster_213 to a band along the rostral medial edge. The former perhaps corresponds to a previously described medial subdivision of the MPul[157]. **(C-D)** We also identified a Group, LGko-IPul Glut, that spans the boundary between the dLG and the inferior pulvinar (IPul). This shared population is consistent with prior microdissection-based transcriptomic studies and immunohistochemistry[45,158], suggesting potential convergent connectivity or shared functions across these nuclei.

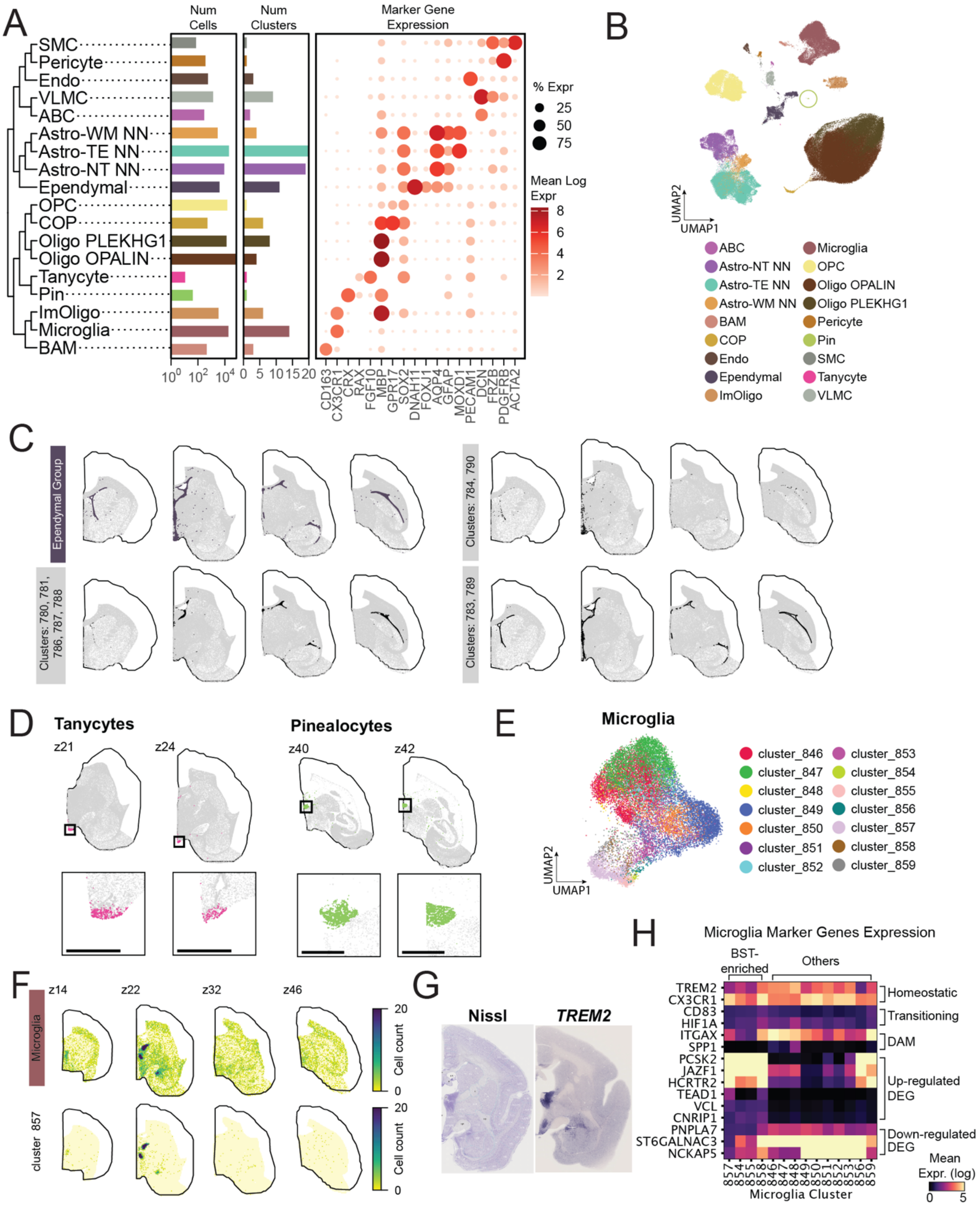


**S5. Spatial localization of non-neuron Groups and related marker gene expression**, related to Fig 1. **(A)** Hierarchically clustered tree of 18 non-neuronal Groups based on the expression of top 4000 HVGs (left). Bar plots indicate cell and Cluster counts per Group (middle). Dot plot shows the mean log expression of canonical marker gene expression for each non-neuron Group (right). **(B)** UMAP of non-neurons colored by Group. **(C)** 2D histograms of cell counts for the ependymal Group (top) and three clusters with localization near specific ventricles (bottom). Cluster_780-781, 786-788 are localized to the posterior lateral ventricles and are likely choroid plexus ependymal types. Cluster_784 and 790 showed strong regional bias to the third ventricle (3V) while Cluster_783 and 789 spanned all ventricles. **(D)** Spatial localization of tanycytes (Cluster_840) (left) to the 3V and pinealocyte Group (Cluster_1854) to the pineal gland (right). **(E)** UMAP of 18,369 Microglia colored by Cluster identity (n=14). **(F)** 2D histogram of cell counts for the Microglia Group indicated the most prominent increase in microglial density in the BNST and HTH (top). Cluster_857 (in addition to Cluster_854-855, 858) drives the high density of microglia in the BST (bottom). Bins for all 2D histograms are 150 um x 150. **(G)** Nissl and *TREM2* ISH staining from the Marmoset Gene Atlas[12,38] (1Y, Male) exhibits increased overall cell density as well as enriched expression of microglia marker *TREM2* in the BST and hypothalamus. **(H)** Marker gene expression across microglial Clusters grouped by whether they were localized to the BST. BST-enriched clusters do not show differential expression of canonical microglial marker genes (*TREM2*, *CX3CR1*) and state-related genes (*CD83*, *HIF1A*, *ITGAX*, *SPP1*). *PCSK2*, *JAZF1* and *HCRTR2* are up-regulated in Cluster 857 when compared against the non-BST enriched Clusters. When comparing all BST-enriched microglia clusters (Clusters 857, 854, 855 and 858) against other microglia clusters as background, top ranked genes include *TEAD1*, *VCL*, *CNRIP1* and negative marker genes include *PNPLA7*, *ST6GALNAC3*, and *NCKAP5*.

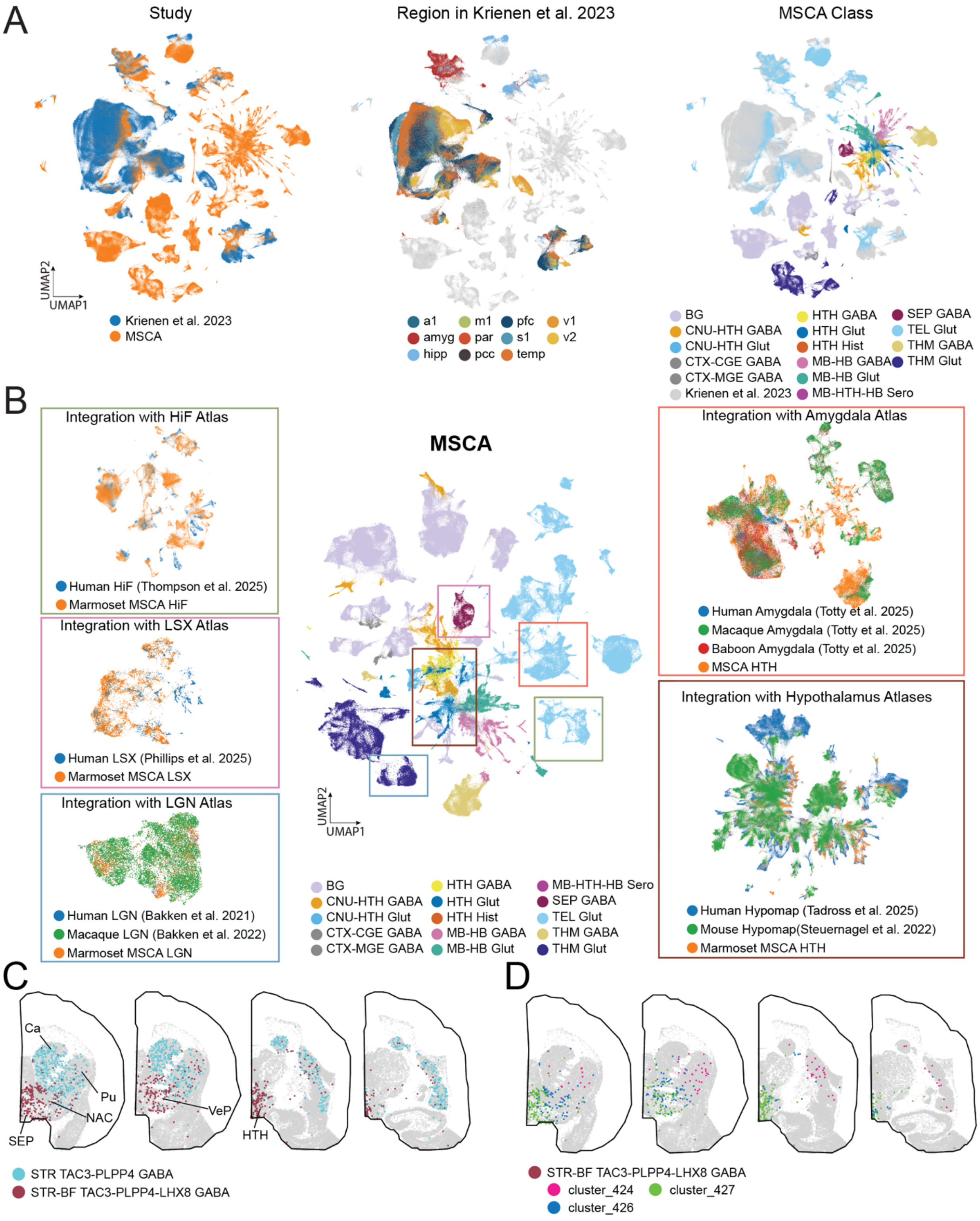


**S6. Integration of MSCA with reference datasets**, related to Fig 1 and Tables 4-5. **(A)** Integration of MSCA TEL Glut Class with previous marmoset census atlas by Krienen et al.[20] , which sampled neocortex at eight different locations, to validate the origin of inadvertent cortical sampling during subcortical dissection. The Tel Glut cells from the MSCA dataset integrate well with the neocortical cells from A1, M1, S1, parietal association cortex (par), prefrontal cortex (pfc), and the temporal pole (temp) in the 2023 census dataset. No MSCA cells integrated with cells dissected from V1 and V2 in the 2023 census dataset. This mismatch is consistent with the expected absence of visual cortical samples in the MSCA dataset because we did not sample the caudal-most slabs that contained only cortex. **(B)** Integration of MSCA with regional reference atlases from different species. Center UMAP shows all MSCA neurons colored by Class and boxes approximating which MSCA cells are integrated with the respective regional reference datasets. Each integration UMAPs surrounding are colored by dataset identities to show reference dataset well-mixed with corresponding MSCA cells. Summarized predicted labels from reference datasets onto MSCA cells are available at **Tables S4-5**. **(C)** Spatial mapping of *TAC3*+ GABAergic interneurons previously characterized in the primate striatum[18,32] indicates that one of the Groups (STR-BF TAC3-PLPP4-LHX8 GABA) is widely distributed across the striatum (Ca, Pu, NAC), septum (SEP), ventral pallidum (VeP) and HTH. (D) Clusters within Group STR-BF TAC3-PLPP4-LHX8 GABA show further regional compartmentalization within these structures.

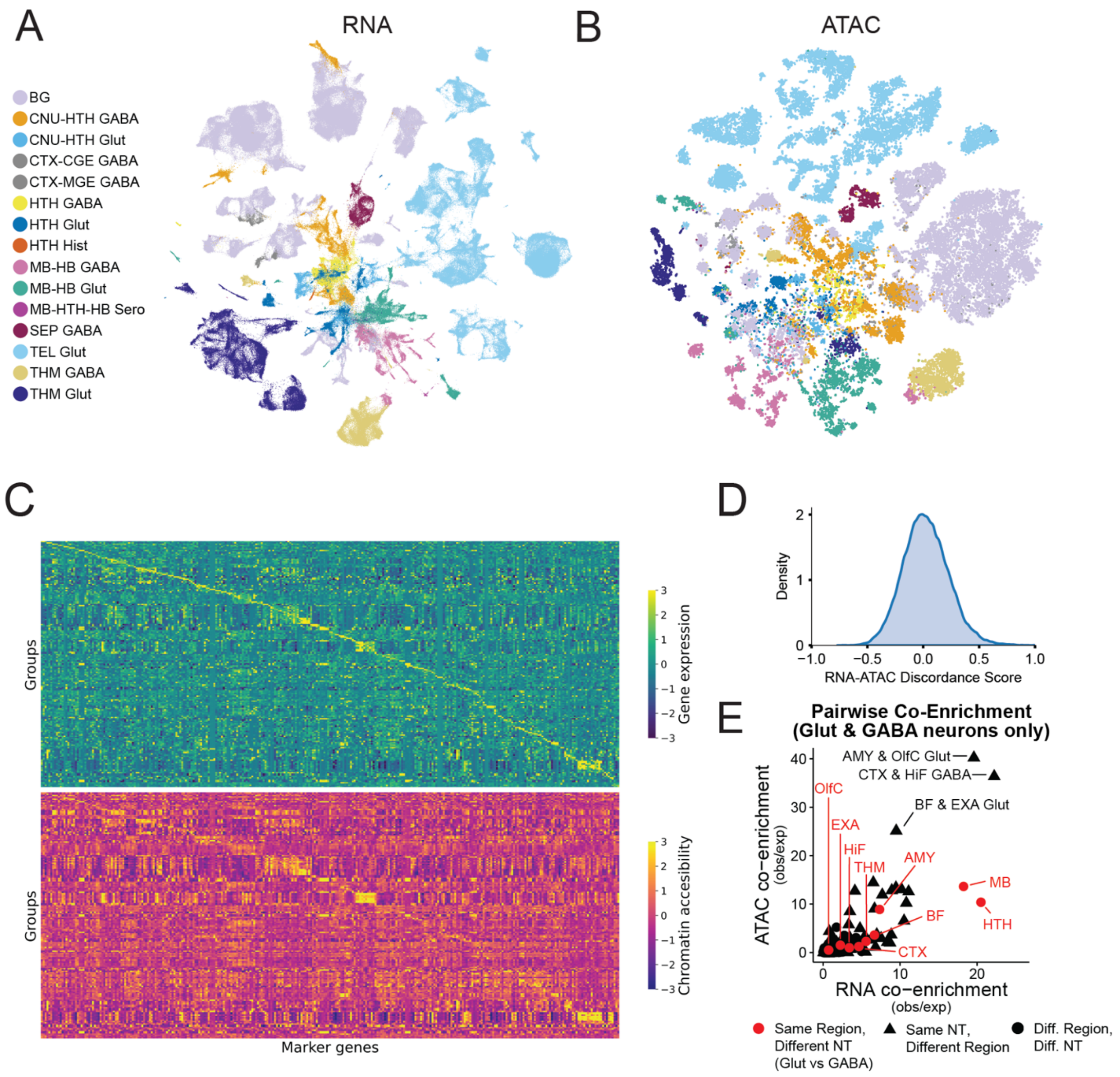


**S7. Concordance between RNA-seq and ATAC-seq data**, related to Fig 1**. (A)** UMAP of all neurons based on gene expression colored by Class. **(B)** tSNE embedding of sub-sampled neurons (n=500000) based on chromatin accessibility-derived cisTopics colored by Class. **(C)** Heatmap of marker gene expression (top) and matched chromatin accessibility of peaks in the corresponding marker gene (bottom) of each group. **(D)** Density plot of RNA-ATAC discordance score calculated from pairwise cosine distances between z-scored Group centroids in RNA (scVI embedding) and ATAC (Topic embedding) latent spaces. **(E)** Neurons are grouped by anatomical region and neurotransmitter. For every possible pair, co-enrichment is based on the number overlaps in the top 1% of genes (RNA) or top 2% of ATAC peaks according to apeglm-shrunken log2FoldChange estimates (see Methods section on co-enrichment analysis).

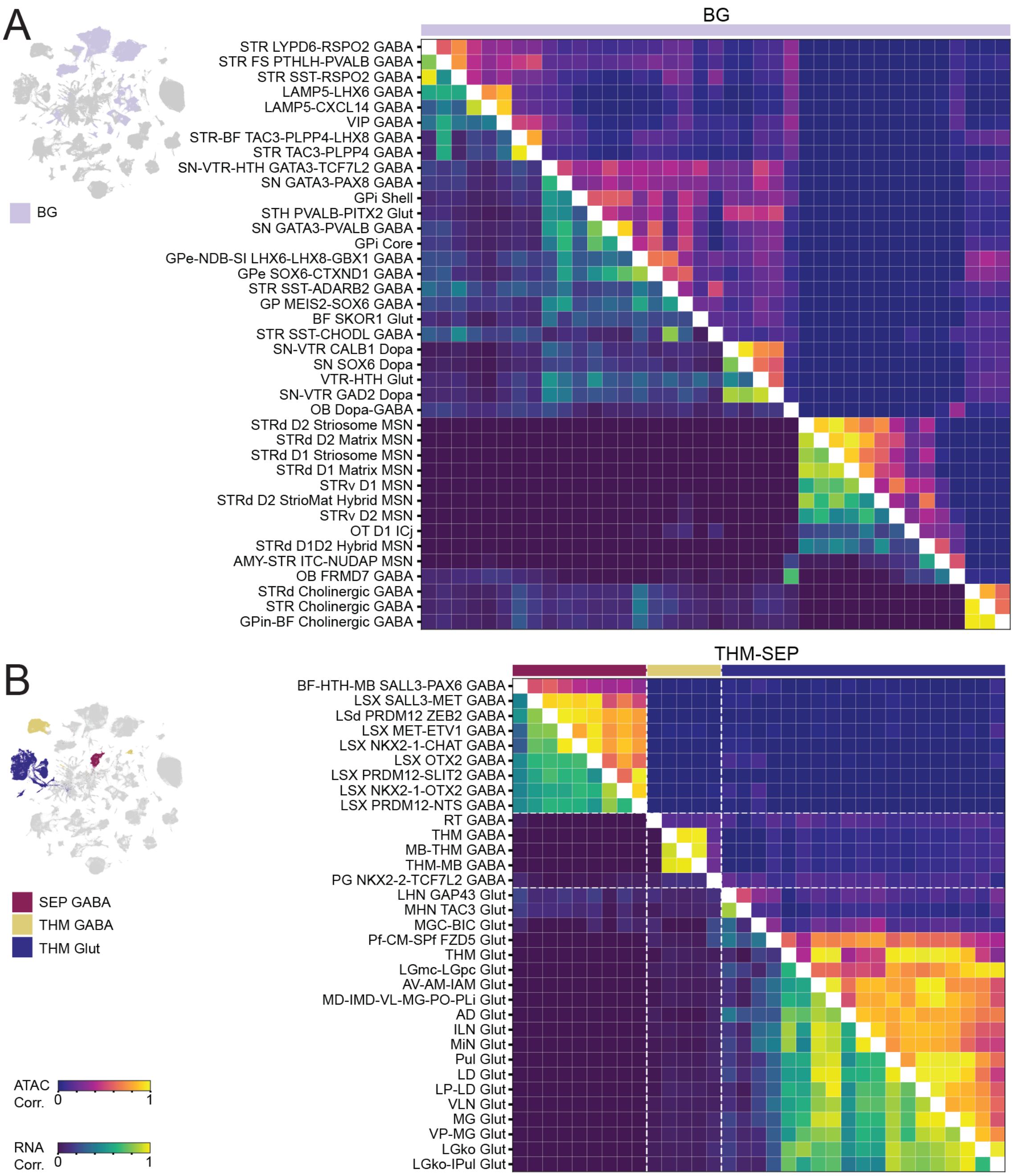

A
BG
BG
STR LYPD6-RSPO2 GABA
STR FS PTHLH-PVALB GABA
STR SST-RSPO2 GABA
LAMP5-LHX6 GABA
LAMP5-CXCL14 GABA
VIP GABA
STR-BF TAC3-PLPP4-LHX8 GABA
STR TAC3-PLPP4 GABA
SN-VTR-HTH GATA3-TCF7L2 GABA
SN GATA3-PAX8 GABA
GPi Shell
STH PVALB-PITX2 Glut
SN GATA3-PVALB GABA
GPi Core
GPe-NDB-SI LHX6-LHX8-GBX1 GABA
GPe SOX6-CTXND1 GABA
STR SST-ADARB2 GABA
GP MEIS2-SOX6 GABA
BF SKOR1 Glut
STR SST-CHODL GABA
SN-VTR CALB1 Dopa
SN SOX6 Dopa
VTR-HTH Glut
SN-VTR GAD2 Dopa
OB Dopa-GABA
STRd D2 Striosome MSN
STRd D2 Matrix MSN
STRd D1 Striosome MSN
STRd D1 Matrix MSN
STRv D1 MSN
STRd D2 StrioMat Hybrid MSN
STRv D2 MSN
OT D1 ICj
STRd D1D2 Hybrid MSN
AMY-STR ITC-NUDAP MSN
OB FRMD7 GABA
STRd Cholinergic GABA
STR Cholinergic GABA
GPin-BF Cholinergic GABA
B
THM-SEP
SEP GABA
THM GABA
THM Glut
BF-HTH-MB SALL3-PAX6 GABA
LSX SALL3-MET GABA
LSd PRDM12 ZEB2 GABA
LSX MET-ETV1 GABA
LSX NKX2-1-CHAT GABA
LSX OTX2 GABA
LSX PRDM12-SLIT2 GABA
LSX NKX2-1-OTX2 GABA
LSX PRDM12-NTS GABA
RT GABA
THM GABA
MB-THM GABA
THM-MB GABA
PG NKX2-2-TCF7L2 GABA
LHN GAP43 Glut
MHN TAC3 Glut
MGC-BIC Glut
Pf-CM-SPf FZD5 Glut
THM Glut
LGmc-LGpc Glut
AV-AM-IAM Glut
MD-IMD-VL-MG-PO-PLi Glut
AD Glut
ILN Glut
MiN Glut
Pul Glut
LD Glut
LP-LD Glut
VLN Glut
MG Glut
VP-MG Glut
LGko Glut
LGko-IPul Glut
ATAC Corr. 0 1
RNA Corr. 0 1

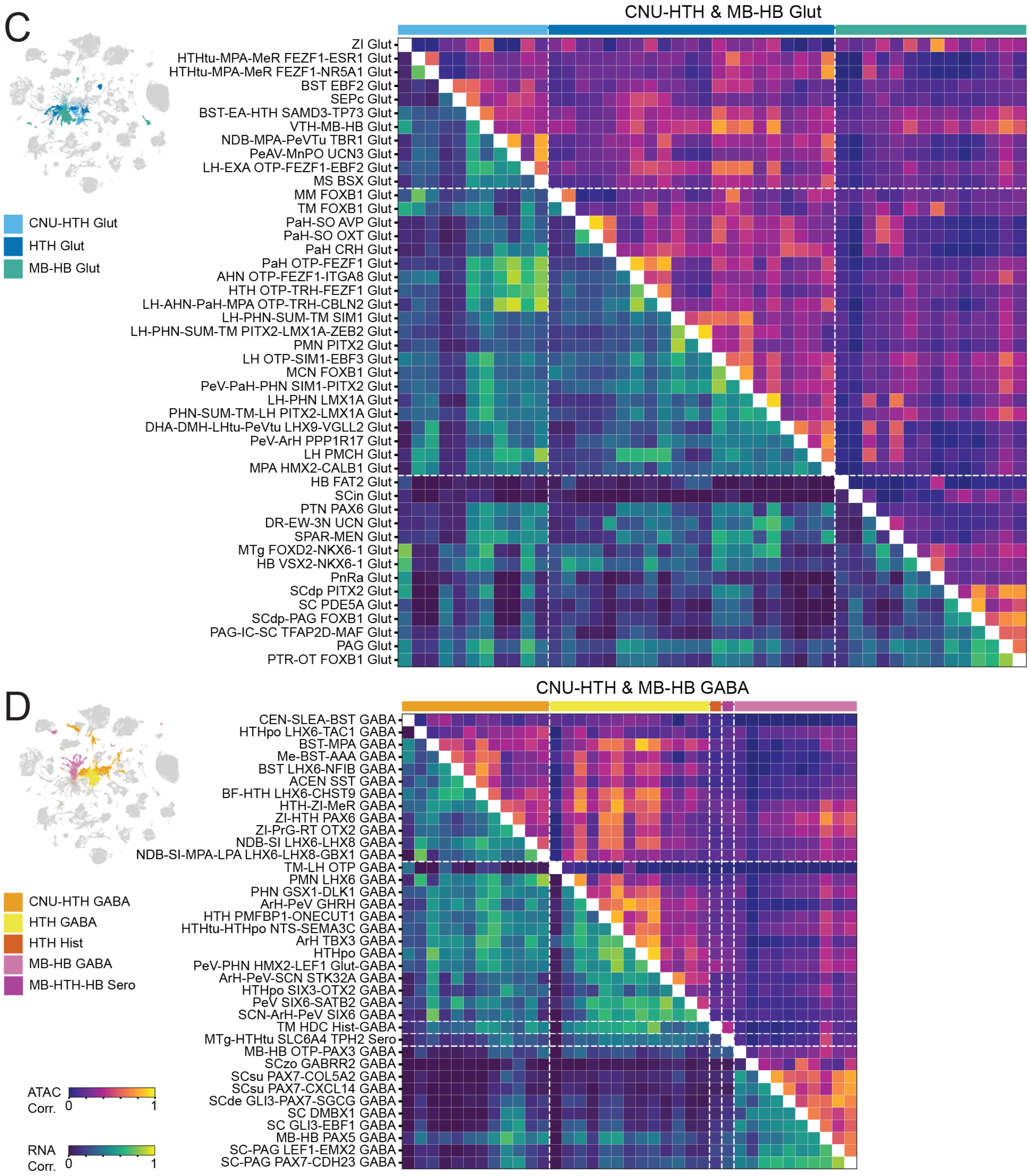
C
CNU-HTH & MB-HB Glut
CNU-HTH Glut
HTH Glut
MB-HB Glut
ZI Glut
HTHtu-MPA-MeR FEZF1-ESR1 Glut
HTHtu-MPA-MeR FEZF1-NR5A1 Glut
BST EBF2 Glut
SEPc Glut
BST-EA-HTH SAMD3-TP73 Glut
VTH-MB-HB Glut
NDB-MPA-PeVTu TBR1 Glut
PeAV-MnPO UCN3 Glut
LH-EXA OTP-FEZF1-EBF2 Glut
MS BSX Glut
MM FOXB1 Glut
TM FOXB1 Glut
PaH-SO AVP Glut
PaH-SO OXT Glut
PaH CRH Glut
PaH OTP-FEZF1 Glut
AHN OTP-FEZF1-ITGA8 Glut
HTH OTP-TRH-FEZF1 Glut
LH-AHN-PaH-MPA OTP-TRH-CBLN2 Glut
LH-PHN-SUM-TM SIM1 Glut
LH-PHN-SUM-TM PITX2-LMX1A-ZEB2 Glut
PMN PITX2 Glut
LH OTP-SIM1-EBF3 Glut
MCN FOXB1 Glut
PeV-PaH-PHN SIM1-PITX2 Glut
LH-PHN LMX1A Glut
PHN-SUM-TM-LH PITX2-LMX1A Glut
DHA-DMH-LHtu-PeVtu LHX9-VGLL2 Glut
PeV-ArH PPP1R17 Glut
LH PMCH Glut
MPA HMX2-CALB1 Glut
HB FAT2 Glut
SCin Glut
PTN PAX6 Glut
DR-EW-3N UCN Glut
SPAR-MEN Glut
MTg FOXD2-NKX6-1 Glut
HB VSX2-NKX6-1 Glut
PnRa Glut
SCdp PITX2 Glut
SC PDE5A Glut
SCdp-PAG FOXB1 Glut
PAG-IC-SC TFAP2D-MAF Glut
PAG Glut
PTR-OT FOXB1 Glut
D
CNU-HTH & MB-HB GABA
CNU-HTH GABA
HTH GABA
HTH Hist
MB-HB GABA
MB-HTH-HB Sero
ATAC Corr. 0 1
RNA Corr. 0 1
CEN-SLEA-BST GABA
HTHpo LHX6-TAC1 GABA
BST-MPA GABA
Me-BST-AAA GABA
BST LHX6-NFIB GABA
ACEN SST GABA
BF-HTH LHX6-CHST9 GABA
HTH-ZI-MeR GABA
ZI-HTH PAX6 GABA
ZI-PrG-RT OTX2 GABA
NDB-SI LHX6-LHX8 GABA
NDB-SI-MPA-LPA LHX6-LHX8-GBX1 GABA
TM-LH OTP GABA
PMN LHX6 GABA
PHN GSX1-DLK1 GABA
HTH PMFBP1-ONECUT1 GABA
ArH-PeV GHRH GABA
HTHtu-HTHpo NTS-SEMA3C GABA
ArH TBX3 GABA
HTHpo GABA
PeV-PHN HMX2-LEF1 Glut-GABA
ArH-PeV-SCN STK32A GABA
HTHpo SIX3-OTX2 GABA
PeV SIX6-SATB2 GABA
SCN-ArH-PeV SIX6 GABA
TM HDC Hist-GABA
MTg-HTHtu SLC6A4 TPH2 Sero
MB-HB OTP-PAX3 GABA
SCzo GABRR2 GABA
SCsu PAX7-COL5A2 GABA
SCsu PAX7-CXCL14 GABA
SCde GLI3-PAX7-SGCG GABA
SC DMBX1 GABA
SC GLI3-EBF1 GABA
MB-HB PAX5 GABA
SC-PAG LEF1-EMX2 GABA
SC-PAG PAX7-CDH23 GABA

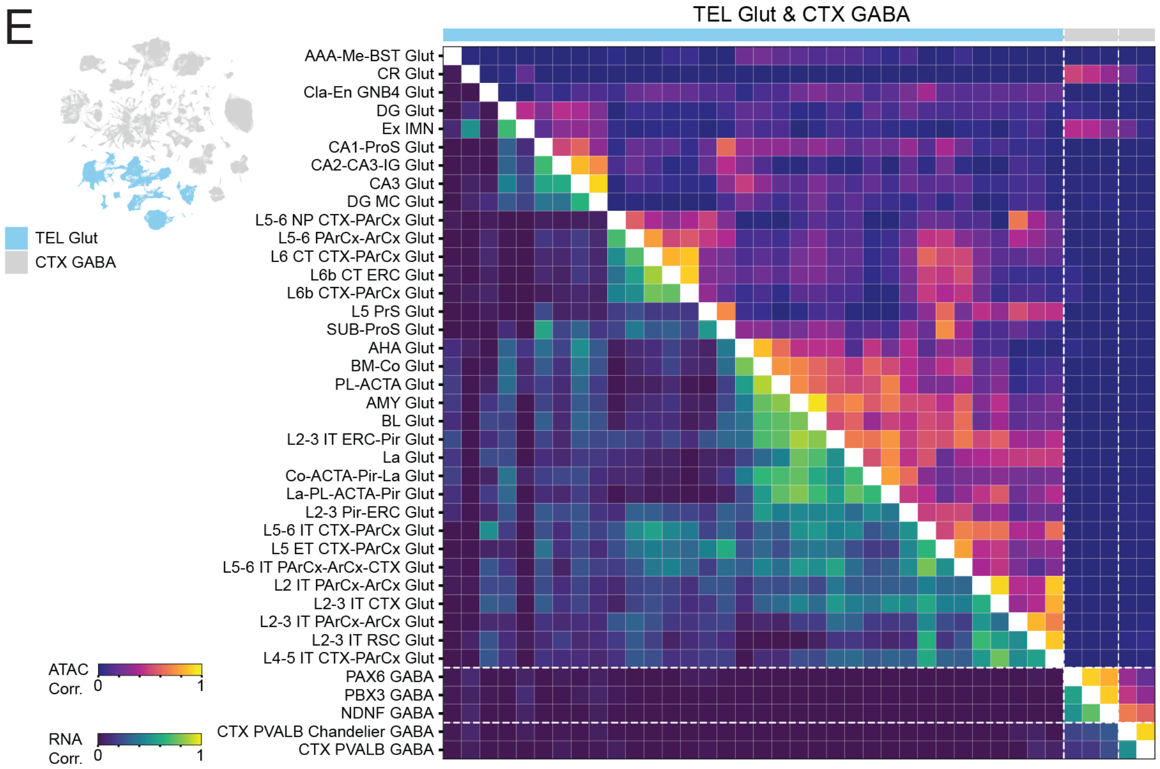


**S8. Pairwise correlation of transcriptomic and chromatin accessibility similarity between Groups within selected Classes,** related to Fig 1. Heatmaps (right) showing pairwise correlation of each group in Class(es) **(A)** BG, **(B)** THM-SEP (THM GABA, THM Glut, SEP-GABA) **(C)** CNU-HTH & MB-HB Glut (CNU-HTH Glut, HTH Glut, MB-HB Glut), **(D)** CNU-HTH & MB-HB GABA (CNU-HTH GABA, HTH GABA, HTH Hist, MB-HTH-HB Sero, MB-HB GABA) **(E)** Tel Glut & CTX GABA (CTX-CGE GABA, CTX-MGE GABA). In each heatmap, rows and columns represent the same Groups, ordered identically as labeled on the y-axis. Upper-left triangles show ATAC correlation and lower-right triangles show RNA correlation between Group-level profiles, scaled from 0 to 1. Colored bars on top of the heatmaps indicate the Class identity of a given Group. UMAP (left) are colored by Class(es) shown in each corresponding heatmap.

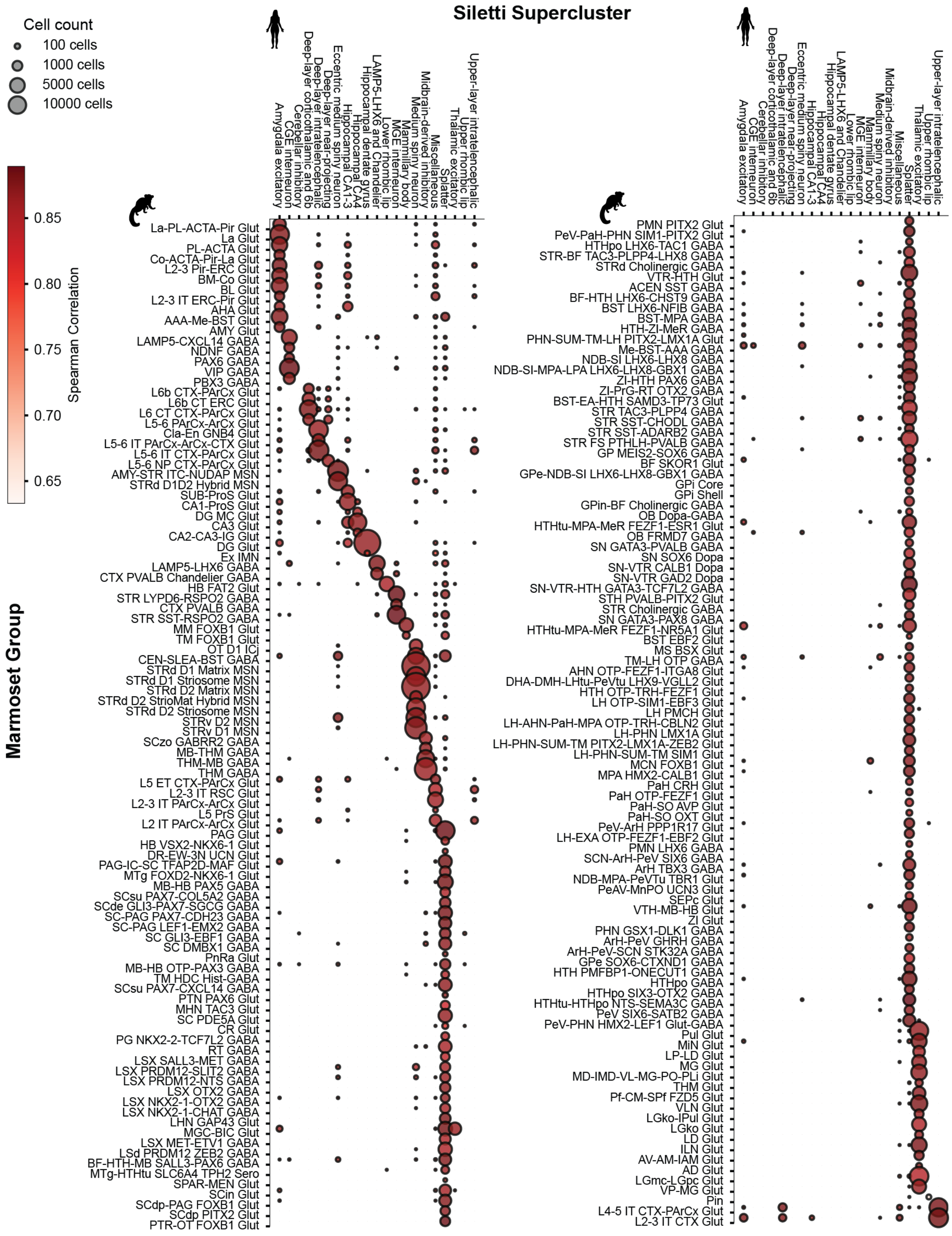


**S9. MapMyCells predictions of Supercluster from the human brain atlas[3] on marmoset cells aggregated by Group**, related to Fig 2. Dot size represents the number of cells and dot color shows the spearman correlation of gene expression between the corresponding marmoset Groups and human Superclusters.

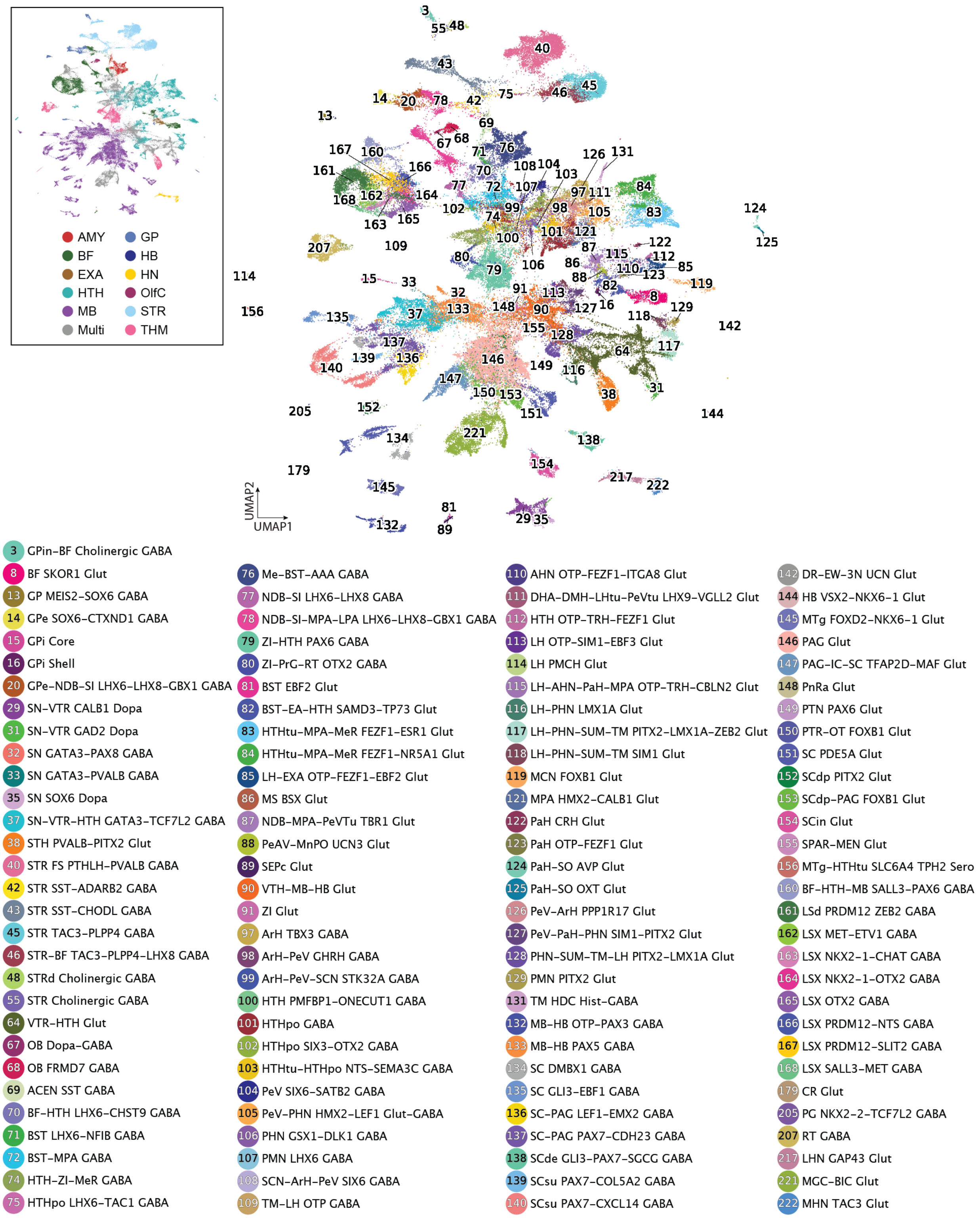


**S10. UMAP embedding of all MSCA cells mapped to human Splatter Supercluster colored by MSCA Groups discussed in the manuscript**, related to Fig 2.

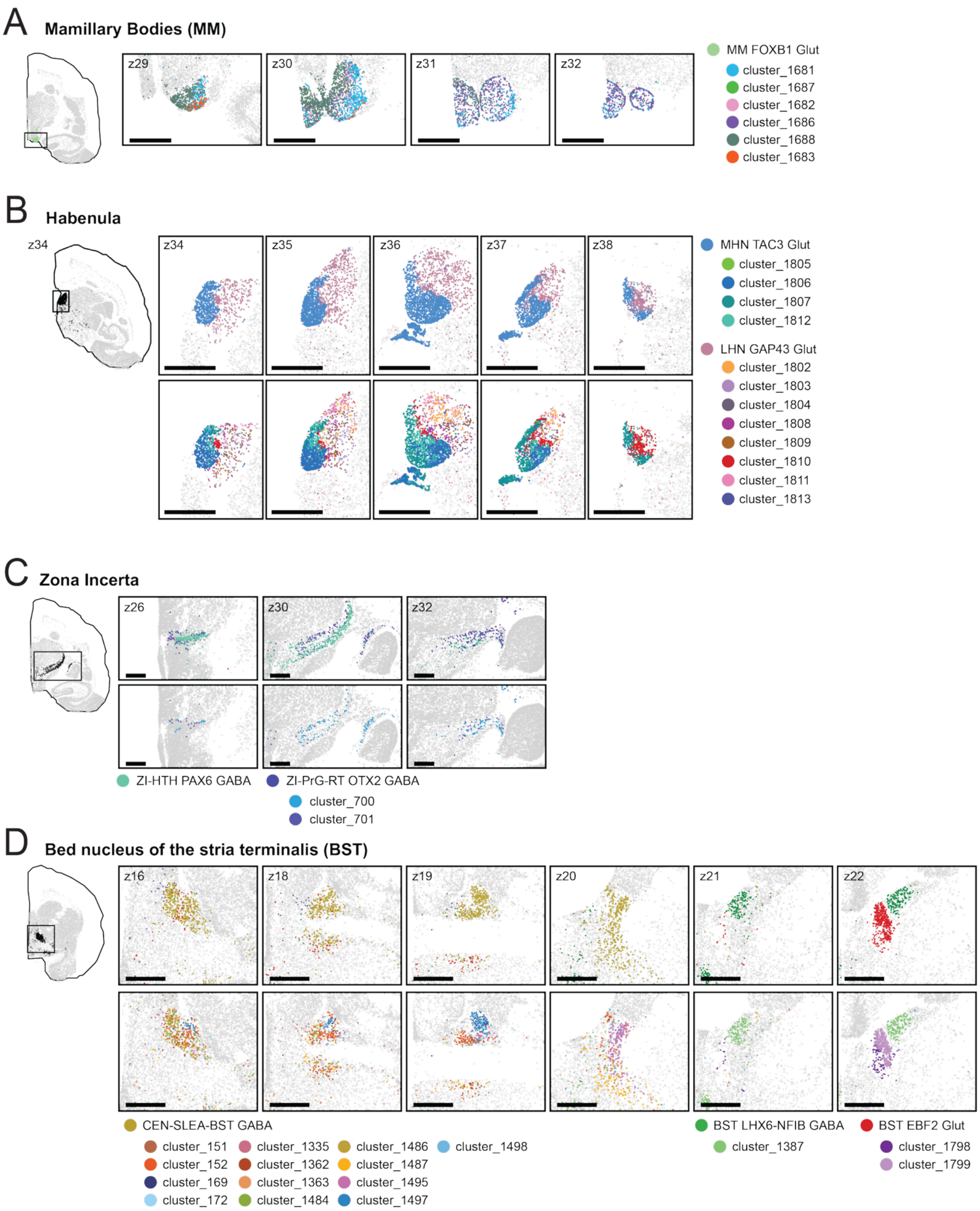


**S11. Representative spatial sections of "Splatter" Groups at the Cluster level show more intricate spatial patterns in diverse brain regions**, related to Fig 2. **(A)** Mammillary bodies **(B)** Habenula **(C)** Zona Incerta **(D)** Bed nucleus of the stria terminalis

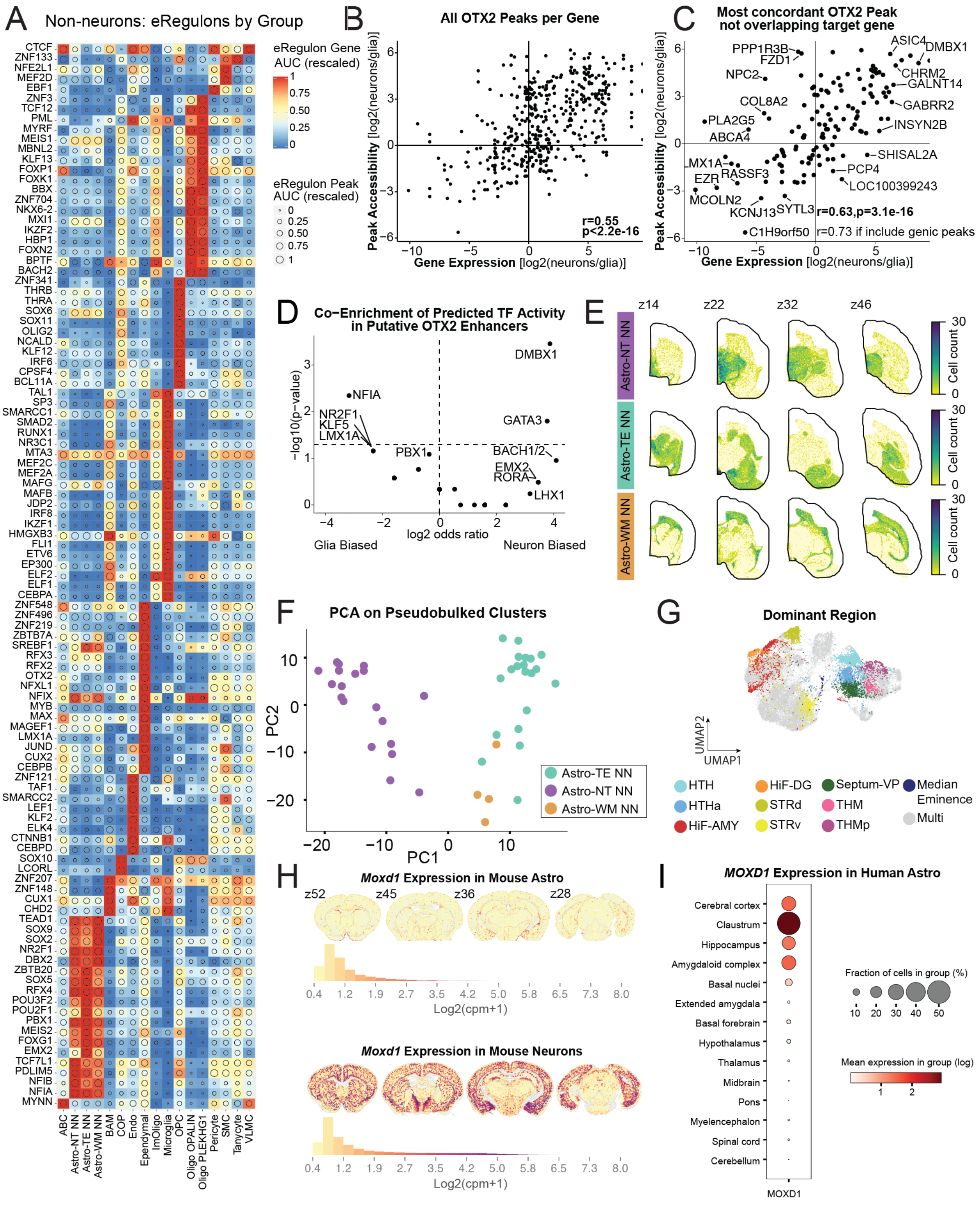


**S12. Non-neuronal activator eRegulons and regional zonation of astrocytes**, related to Fig 3 and Fig 4. **(A)** Heatmap showing all direct activator (+/+) eRegulons detected in non-neurons. Non-neurons grouped at the Group level. **(B)** Comparison of OTX2 target gene expression and the accessibility of peak with inferred OTX2 binding/activity. Of 436 peaks, 202 (46%) are within the target gene itself (90 within 500bp of any TSS, and 57 peaks contain the target gene TSS). **(C)** Same as B, but peaks were subset to only those not found within the target gene. Of those, the peak with most similar enrichment to the target gene's expression was selected. **(D)** Peaks with predicted OTX2 activity sometimes have other TFs with predicted activity. Each point represents a candidate co-binding TF detected in the same OTX2-bound peaks linked to differentially expressed (in neurons vs. non-neurons) target genes. Peaks were selected if both they and their target genes were neuron- or nonneuron-biased (log2FoldChange magnitude >0.75), which produces a set of n=198 selected OTX2 peaks (associated with 130 unique gene targets). Odds ratio is for co-binding in those selected peaks, and p-values are unadjusted from Fisher's exact test (dashed line indicates p=0.05). **(E)** 2D histogram of cell counts showing the regional zonation of the three astrocyte Groups. **(F)** UMAP embedding of all astrocytes colored by their dominant region. **(G)** Pseudobulked Cluster expression profiles in PC space. Each dot is an astrocyte Cluster and colored by their Group identity. **(H)** *Moxd1* expression in mouse astrocytes (top) and mouse neurons (bottom) (Allen Brain Cell Atlas, RRID: SCR_024440). **(I)** *MOXD1* expression in human astrocytes grouped by dissection regions[3].

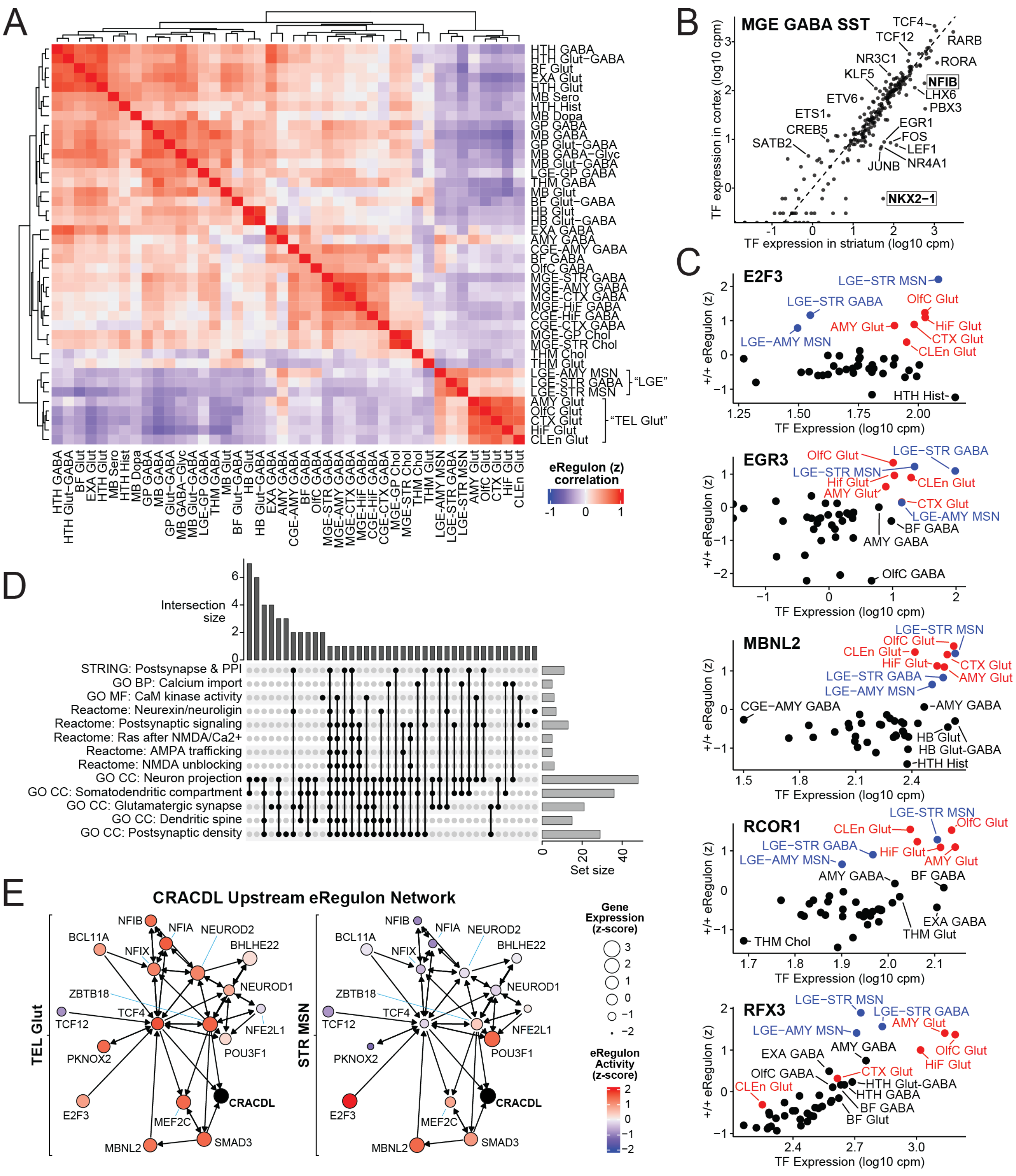


**S13. Extended results on the shared regulatory programs in striatal MSNs and telencephalic glutamatergic neurons**, related to Fig 5. **(A)** As in Fig 5A,E: Cluster-level eRegulon scores (z-score of gene AUC) are averaged by region and neurotransmitter, with MSNs and GE-derived clusters separated. Pearson correlations are shown. **(B)** As in **Fig5D**, but for the SST subtype of MGE GABA. **(C)** For selected eRegulons co-enriched in TEL Glut & LGE GABA, the TF's expression and eRegulon (direct +/+) activity (gene AUC z-score) are shown. Both values are means for all clusters in each grouping. **(D)**

Related to the string-db enrichment analysis of genes co-enriched in TEL Glut & LGE GABA, “Upset” plot shows selected terms for co-enriched genes. BP = Gene Ontology (GO) Biological Process; MF = Molecular Function; CC = Cellular Component. **(E)** All direct eRegulons within two hops of CRACDL are shown, with nodes colored by gene AUC z-scores and sized by gene expression z-scores.

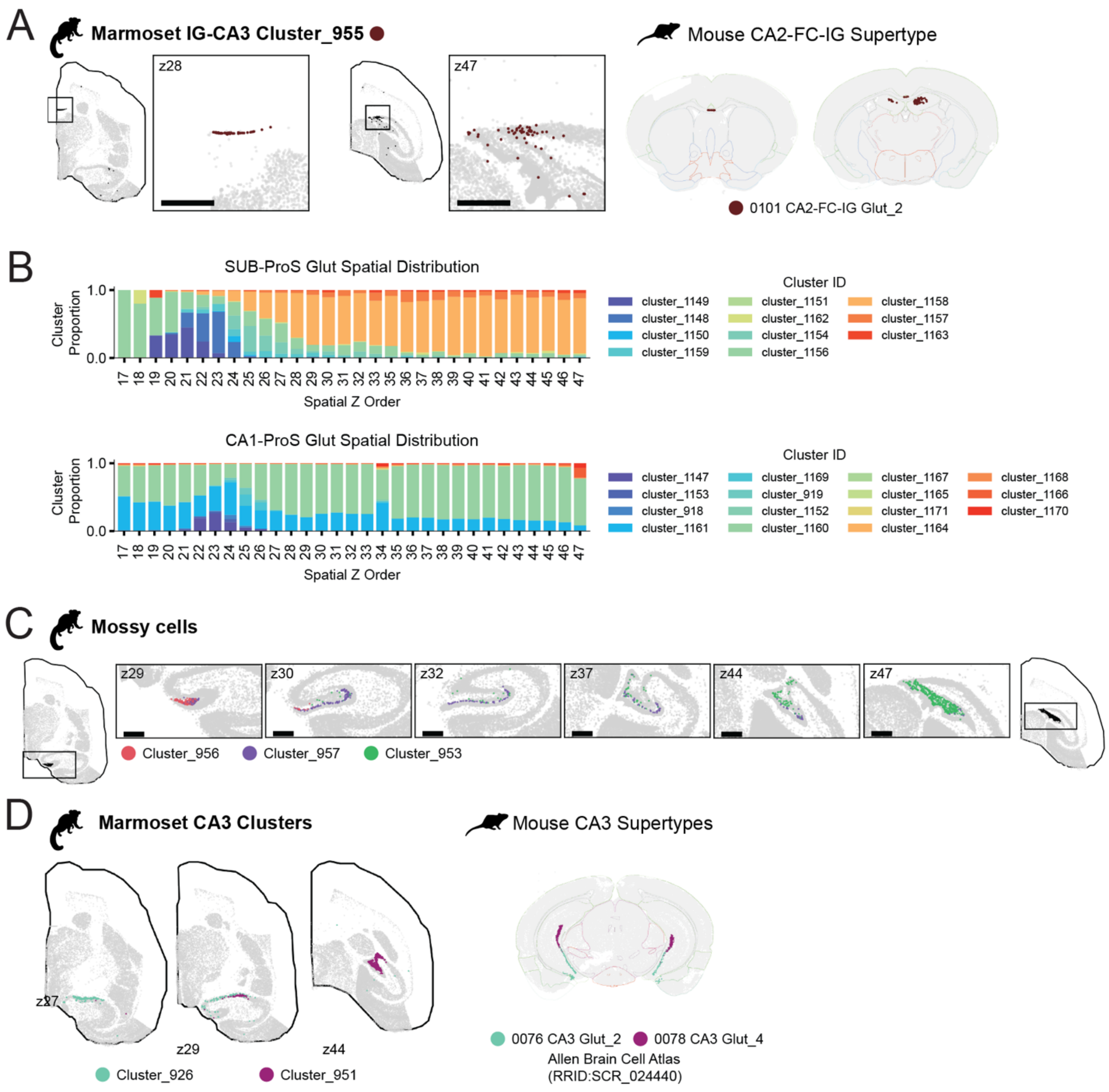


**S14. Spatial distribution of transcriptomic clusters in the hippocampus**, related to Fig 6. **(A)** Marmoset cluster 995 localizes to IG and posterior CA3 (left), and maps to mouse supertype 0101 CA2-FC-IG Glut_2 (right). **(B)** Cluster proportion changes from anterior to posterior HiF in SUB-ProS Glut (top) and CA1-ProS Glut (bottom). **(C)** Spatial localization of mossy cells Clusters in DG with Cluster_956 restricted to anterior sections and Cluster_953 localized to posterior sections. **(D)** Clusters with anterior or posterior bias in marmoset CA3 (left) map to mouse ventral and dorsal CA3 supertypes (right), respectively.

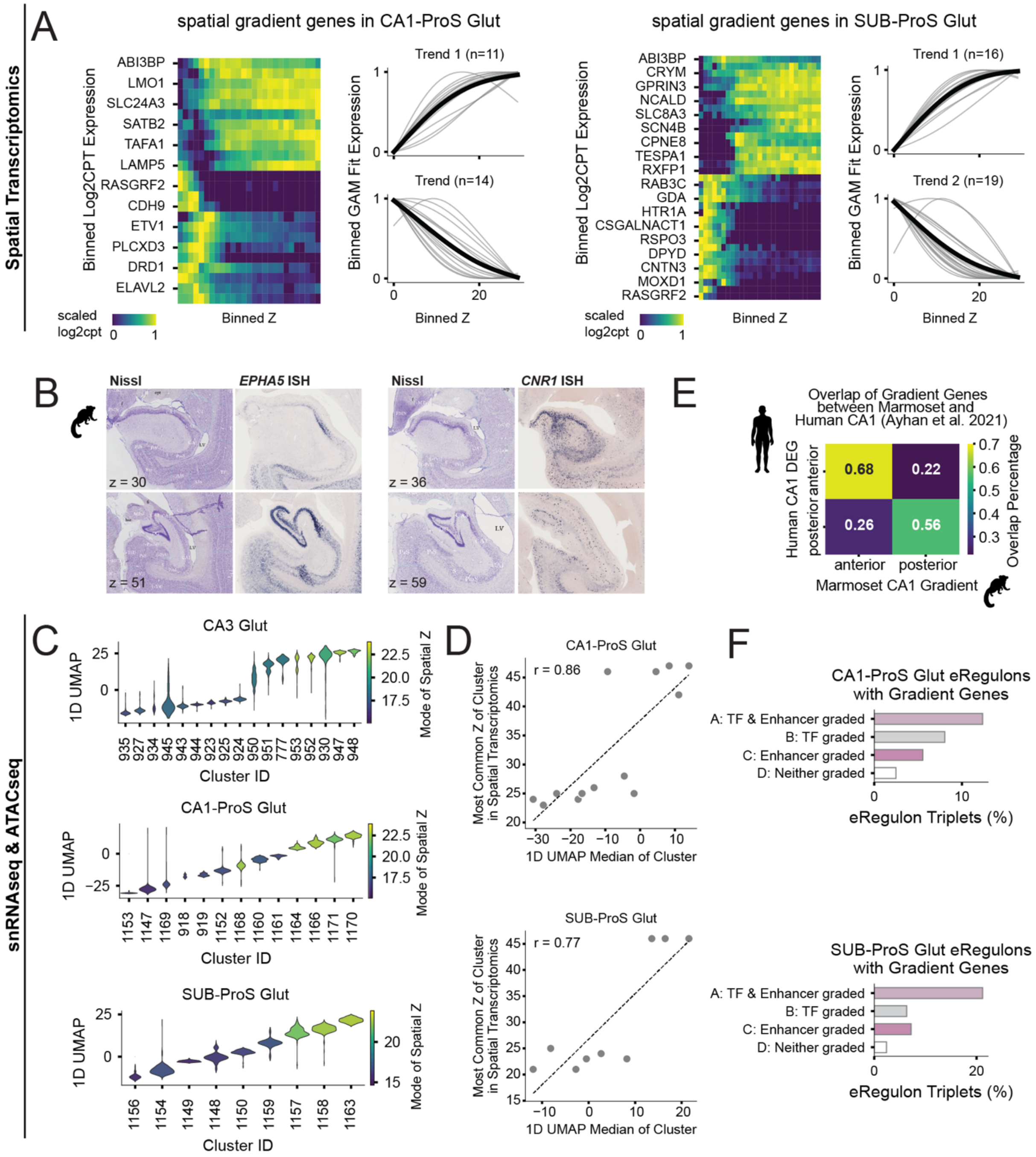


**S15. Extended results on spatial gene gradients and regulatory programs in the hippocampus**, related to Fig 6. **(A)** Left: Log2cpt expression of spatial gradient genes found in the corresponding Group (CA1-ProS Glut, SUB-ProS Glut) in each spatial bin (n=30) ordered anterior to posterior from left to right. Rows are ordered based on the hierarchical clustered GAM fit. Right: GAM-fitted expression of genes across spatial bins and separated into 2 major trends. The number of genes assigned to each trend is shown in parentheses. In both Groups, the majority of genes have increased expression posteriorly in trend 1 while genes in trend 2 are characterized by decreasing posterior expression. **(B)** Left: *EPHA5* ISH staining from the Marmoset Gene Atlas showing higher expression of *EPHA5* in posterior CA3 in comparison to anterior CA3 (3M, male). Right: *CNR1* ISH staining from the Marmoset Gene Atlas[12,38]

showing higher expression of *CNR1* in posterior CA1-ProS in comparison to anterior CA1-ProS (1Y, Female). **(C)** Distribution of 1D UMAP coordinates for cells in each Cluster of CA3 Glut (top), CA1-ProS Glut (middle) and SUB-ProS Glut (bottom) Group, with color indicating the modal spatial z-order of the cluster. **(D)** Scatter plots showing correlation of median 1D UMAP coordinates and modal spatial z-section for each Cluster in CA1-ProS Glut (top; r=0.86) and SUB-ProS Glut (bottom; r=0.77). **(E)** Heatmap of percentage of marmoset CA1 gradient genes (p-adj<0.05) that overlap with differentially expressed genes in anterior and posterior human CA1[109]. **(F)** Percentage of eRegulon triplets with target gene gradients under each possibility outlined in **Fig 6F** in CA1-ProS Glut (top) and SUB-ProS Glut (bottom).

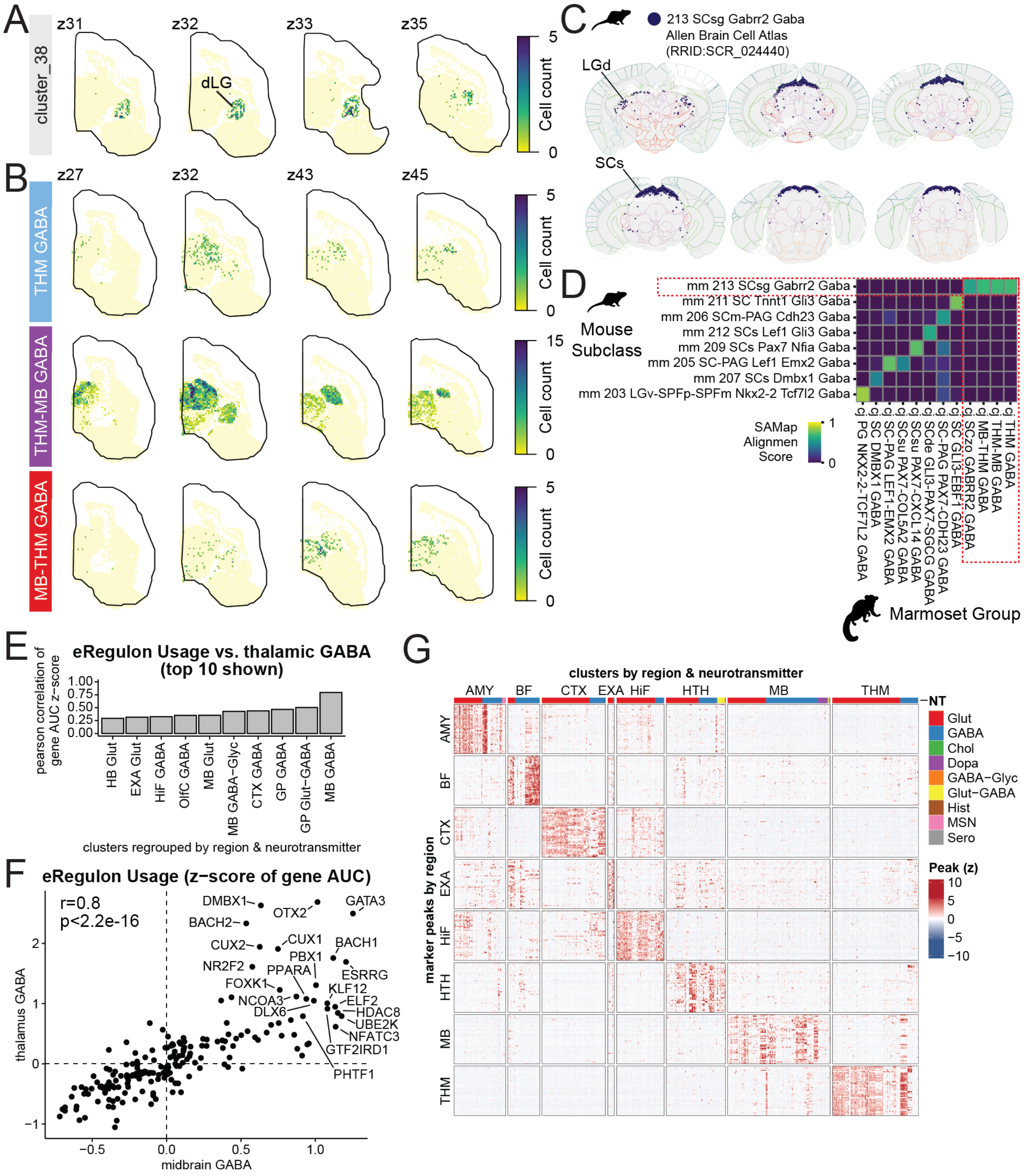


**S16. Spatial localization and regulatory landscape of GABAergic Groups in the primate Thalamus**, related to Fig 7. **(A)** 2D histograms of cell counts showing the regional localization of Cluster_38 to the dLG. **(B)** 2D histograms of cell counts for the three non-RT thalamic GABAergic Groups. **(C)** SAMap cross-species alignment scores of marmoset non-RT thalamic and SC GABAergic Groups against mouse SC GABAergic subclasses. Red dashed outlines highlight the alignment scores between marmoset thalamic GABAergic Groups and mouse subclass 213 SCsg Gabrr2 Gaba. **(D)** Spatial localization of mouse subclass 213 SCsg Gabrr2 Gaba from the Allen Brain Cell Atlas (RRID:SCR_024440). The population predominates in the superior colliculus of the mouse midbrain, though also extends a sparse population to the LGN. **(E)** Top 10 populations (aggregated by dominant region and neurotransmitter identity) correlated with thalamic GABAergic neurons (from Groups THM GABA, THM-MB GABA, RT GABA, ZI-HTH PAX6 GABA (1 Cluster excluded), ZI-PrG-RT OTX2 GABA, PG NKX2-2-TCFL2 GABA) by average eRegulon usage quantified by cluster gene AUC z-scores. **(F)** Specific eRegulon usage (gene AUC z-scores) are shown for all midbrain- or thalamus-localized GABAergic neurons. **(G)** Top regional marker peaks identified in each anatomical region.
